# Distinct involvements of the subthalamic nucleus subpopulations in reward-biased decision-making in monkeys

**DOI:** 10.1101/2025.11.04.686631

**Authors:** Kathryn Branam, Joshua I. Gold, Long Ding

## Abstract

The subthalamic nucleus (STN) is a part of the indirect and hyperdirect pathways in the basal ganglia (BG) and has been implicated in movement control, impulsivity, and decision-making. We recently demonstrated that, for perceptual decisions, the STN includes at least three subpopulations of neurons with different decision-related activity patterns (Branam et al., 2024). Here we show that, for decisions that require both perceptual and reward-based processing, many STN neurons are sensitive to both sensory evidence and reward expectations. Within a drift-diffusion framework, three STN subpopulations show different relationships to model components reflecting formation of the decision variable, dynamics of the decision bound, and non-decision-related processes. Many STN neurons also represent quantities related to decision evaluation, including choice accuracy and reward expectation. These results help to further delineate the multiple roles that STN plays in forming and evaluating complex decisions that combine multiple sources of information.

## Introduction

The subthalamic nucleus (STN) is a part of the indirect and hyperdirect pathways in the basal ganglia (BG). The STN provides major glutamatergic inputs to the output nuclei of the BG, namely the substantia nigra pars reticulata (SNr) and the internal segment of the globus pallidus (GPi). Because of its strategic location in these pathways, the STN has been studied extensively in the context of movement control (e.g., Martin, 1927; Martin and Alcock, 1934; Whittier and Mettler, 1949; Carpenter et al., 1950; Wichmann et al., 1994; DeLong and Wichmann, 2001) and is a primary target site for clinical interventions to improve motor functions (e.g., deep brain stimulation in Parkinson’s disease; DeLong and Wichmann, 2001). Functionally, the STN has been linked to inhibitory control of actions or impulsivity (e.g., Baunez et al., 2001; Desbonnet et al., 2004; Witt et al., 2004; Aron and Poldrack, 2006; Frank et al., 2007; Isoda and Hikosaka, 2008; Forstmann et al., 2012; Schmidt et al., 2013; Pasquereau and Turner, 2017; Bonnevie and Zaghloul, 2019).

In parallel to these studies in the motor domain, more recent studies have implicated the STN in a variety of more cognitive functions. These functions include reward-based decision-making (van Wouwe et al., 2011; Espinosa-Parrilla et al., 2015; Seymour et al., 2016; Zénon et al., 2016; Nougaret et al., 2022; Pagnier et al., 2024), noisy evidence-based decision-making (Bogacz and Gurney, 2007; Ratcliff and Frank, 2012; Wei et al., 2015; Zaehle et al., 2017; Branam et al., 2024), and decision conflict resolution (Lehericy et al., 2004; Aron et al., 2007; Fumagalli et al., 2011; Brittain et al., 2012; Zaghloul et al., 2012; Zavala et al., 2017). To support these diverse functions, the STN may contain subpopulations of neurons with distinct computational roles.

Consistent with this idea, we and others previously identified subpopulations of STN neurons with distinct activity patterns when recorded during performance of particular tasks (Zavala et al., 2017; Branam et al., 2024). For example, we identified three subpopulations (Branam et al., 2024) that align roughly with three different computational models of STN’s contributions to perceptual decision-making (Bogacz and Gurney, 2007; Ratcliff and Frank, 2012; Wei et al., 2015). However, the specific computational roles that these different subpopulations play in decision-making and other cognitive functions remain not well understood. For example, two of the subpopulations had overall activity patterns that were consistent with two different models in which the STN modulated the decision bound (Ratcliff and Frank, 2012; Wei et al., 2015), but the exact nature of this modulation is not known. The other subpopulation’s general activity patterns were consistent with a model of STN mediating evidence accumulation (Bogacz and Gurney, 2007), but it is unclear if and how this activity contributes to how evidence is weighed, biased, or accumulated.

Our previous attempt to distinguish these alternatives using electrical microstimulation was unsuccessful because that manipulation likely affected highly intermingled subpopulations with different functions. Here, we build on our previous work by recording from individual STN neurons in monkeys performing a more complex decision task that requires integrating noisy evidence accumulation with reward preference. We show that this increased computational task demand can provide new insights into the diverse computational roles played by different STN subpopulations in the cognitive control of behavior.

## Results

We used the same two monkeys as in the previous study of STN neurons (Branam et al., 2024). For that study, the monkeys performed a visual motion discrimination decision task by making a saccade at a self-chosen time to indicate their perception of the global motion direction of a random-dot kinematogram (Figure 1A). Here we added an asymmetric-reward version of the task, for which we separately manipulated the noisy evidence (motion direction and strength) and reward context (a larger juice reward for a correct choice associated with one of the two directions). For each trial, the motion strength and direction were chosen randomly from five values and two directions, respectively. In a block of ~55 trials, one choice was paired with a large reward, and the other was paired with a small reward. The choice-reward association (i.e., “reward context”) was alternated between blocks and signaled to the monkeys at block transitions via changes in the colors of the choice targets. The monkeys were rewarded for correct choices only. Both monkeys showed consistent biases toward the large-reward choice (Figure 1B, C). Details of their performance, including variations across sessions and individuals, have been reported in a previous study (Fan et al., 2018).

**Figure 1:**
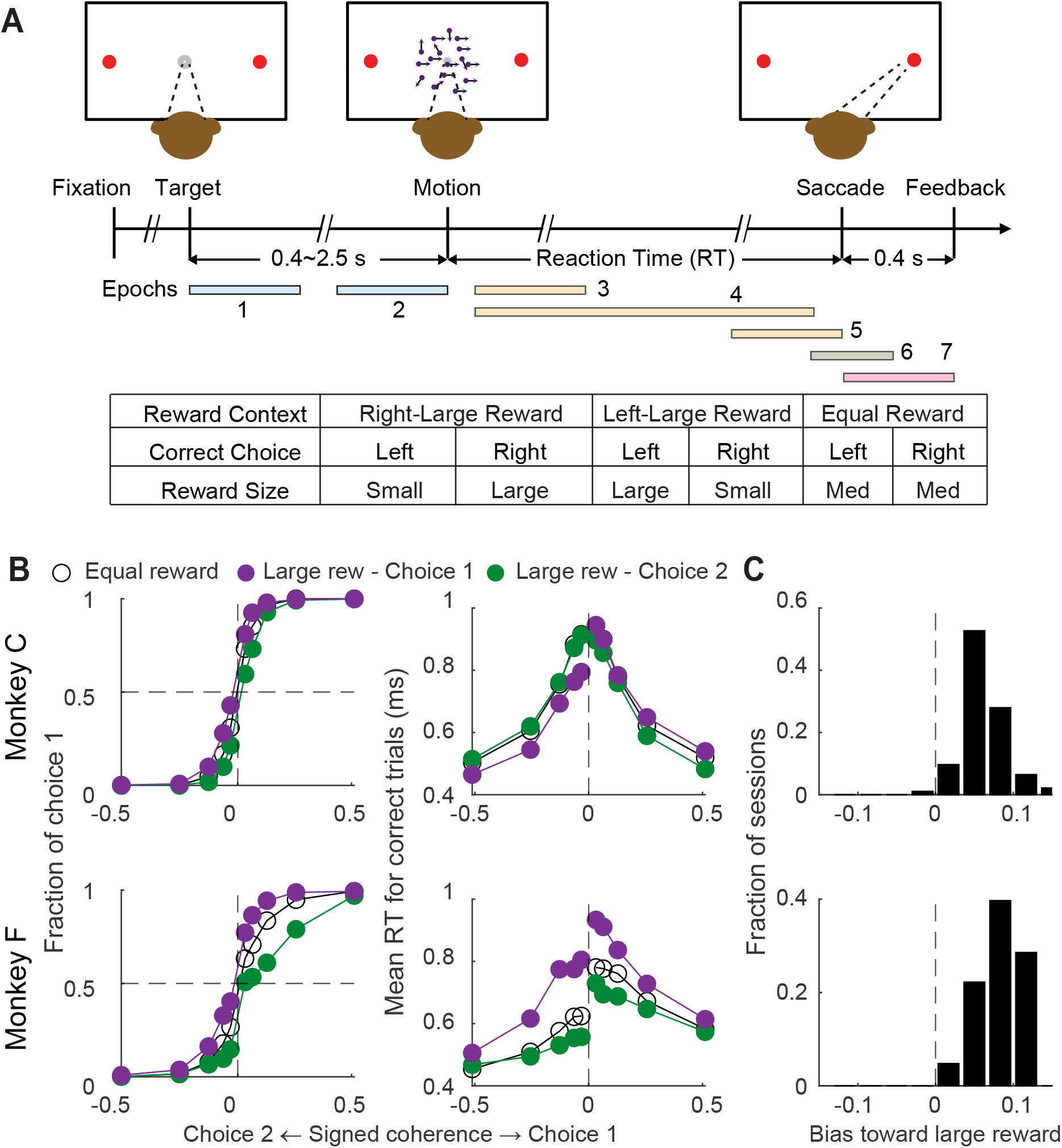
The monkeys were biased toward choices associated with large reward. A, Task design and timeline. The monkeys made saccades to indicate their perceived motion direction. Correct trials were rewarded based on the reward context (see Table inset). Error trials were not rewarded. “Epochs” illustrate the time windows for epoch-based analyses. B, Average choice (left) and RT (right) behavior of the two monkeys. Monkey C: 93 sessions, 37,706 trials; Monkey F: 63 sessions, 27,113 trials. Filled and open circles are data from the three reward contexts, as indicated at the top of the panel. C, Histograms of reward bias for all sessions, estimated using logistic fits to choice data.

### Single STN neurons combine visual and reward information

While the monkeys were performing the equal- and asymmetric-reward versions of the task, we measured single-neuron activity of 156 STN neurons (n=93 and 63 for monkeys C and F, respectively). Many single STN neurons were sensitive to manipulations of visual evidence (the strength and direction of the motion stimulus) and reward contexts (equal rewards, higher reward for the left choice, and higher reward for the right choice). Figure 2 shows the activity of three example neurons on the equal- (panels A–C) and asymmetric- (panels D–F) reward tasks, which highlight the heterogeneity of STN responses, in terms of both the conversion of visual evidence into decisions as previously shown (Branam et al 2024) and interactions between visual evidence and reward information.

**Figure 2:**
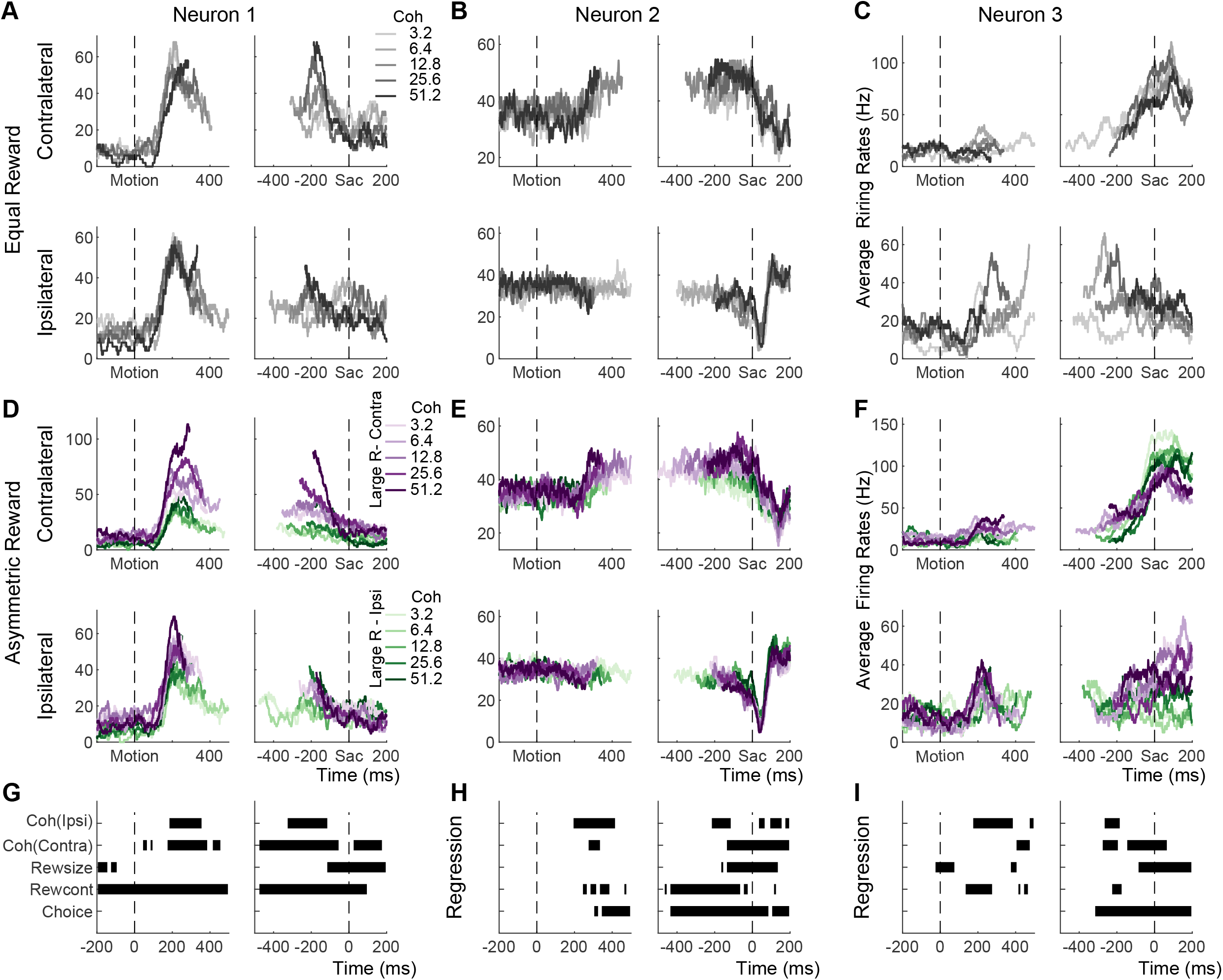
Example STN neurons showing modulation by decision-related factors. A-C, Average activity from the equal-reward task, for trials with contralateral (top row) and ipsilateral (bottom row) choices. Activity was truncated at the median RT for each trial condition. D-F, Average activity from the asymmetric-reward task, for trials with contralateral (top row) and ipsilateral (bottom row) choices. Activity was truncated at the median RT for each trial condition. Purple and green colors indicate blocks with the large reward paired with contralateral and ipsilateral choices, respectively. G-I, Results from a multiple linear regression analysis. Each line shows the timing of significant non-zero coefficients for a specific regressor (t-test, p<0.05). Results for the interaction terms are not shown.

The first example neuron showed choice- and coherence-independent activation after motion onset, which dissipated approaching saccades, on the equal-reward task (Figure 2A). On the asymmetric-reward task, this neuron showed similar activation for the two choices but became sensitive to motion coherence (Figure 2D; e.g., on trials with contralateral choices, dark purple curves representing high-coherence trials are higher than light purple curves representing low-coherence trials). The activation also depended on reward context (purple curves representing trials with higher rewards for right choices are higher than green curves representing trials with higher rewards for left choices). Multiple linear regression results confirmed these visual impressions (Figure 2G).

The second example neuron showed weak choice- and coherence-modulated activation in the late motion viewing period, followed by choice-dependent post-saccade suppression, on the equal-reward task (Figure 2B). On the asymmetric-reward task, the general choice- and coherence-modulation patterns held, with additional reward-context modulation during motion viewing and reward-size modulation around saccade onset (Figure 2E,H; purple curves are higher than green curves for trials with contralateral choices and the reverse for trials with ipsilateral choices).

The third example neuron showed choice-selective activation on the equal-reward task (Figure 2C). On the asymmetric-reward task, this neuron’s activity showed reward-context modulation during early motion viewing, strong choice modulation closer to saccade onset, and reward-size modulation peri- and post-saccades (Figure 2F, I).

Across the population, substantial fractions of STN neurons showed sensitivity to decision-related factors like those evident in the example neurons (Figure 3A). The fraction of neurons with statistically reliable sensitivity for choice (chi-square test, *p*<0.05) increased steadily during motion viewing (epochs indicated by yellow bands) and plateaued around saccade onset. The fraction for reward context stayed above chance level throughout the trial, with the highest values during motion viewing. The fraction for expected reward size rose above chance level during the later stage of motion viewing and persisted through saccades. The fractions for motion coherence were above chance level after motion onset for trials, with similar values for trials with contralateral or ipsilateral choices. There were also small but significant fractions of neurons that were sensitive to the interaction between motion coherence and reward size during motion viewing. Among the neurons with significant modulation, the polarity and timing of modulation varied (Figure 3-S1). Generally, there tended to be more neurons preferring contralateral choices, the context of pairing large rewards with contralateral choices, larger expected reward size, and higher coherences (more purple versus green pixels in the heatmaps). The peak modulation times varied among neurons for all decision-related factors and both polarities.

**Figure 3:**
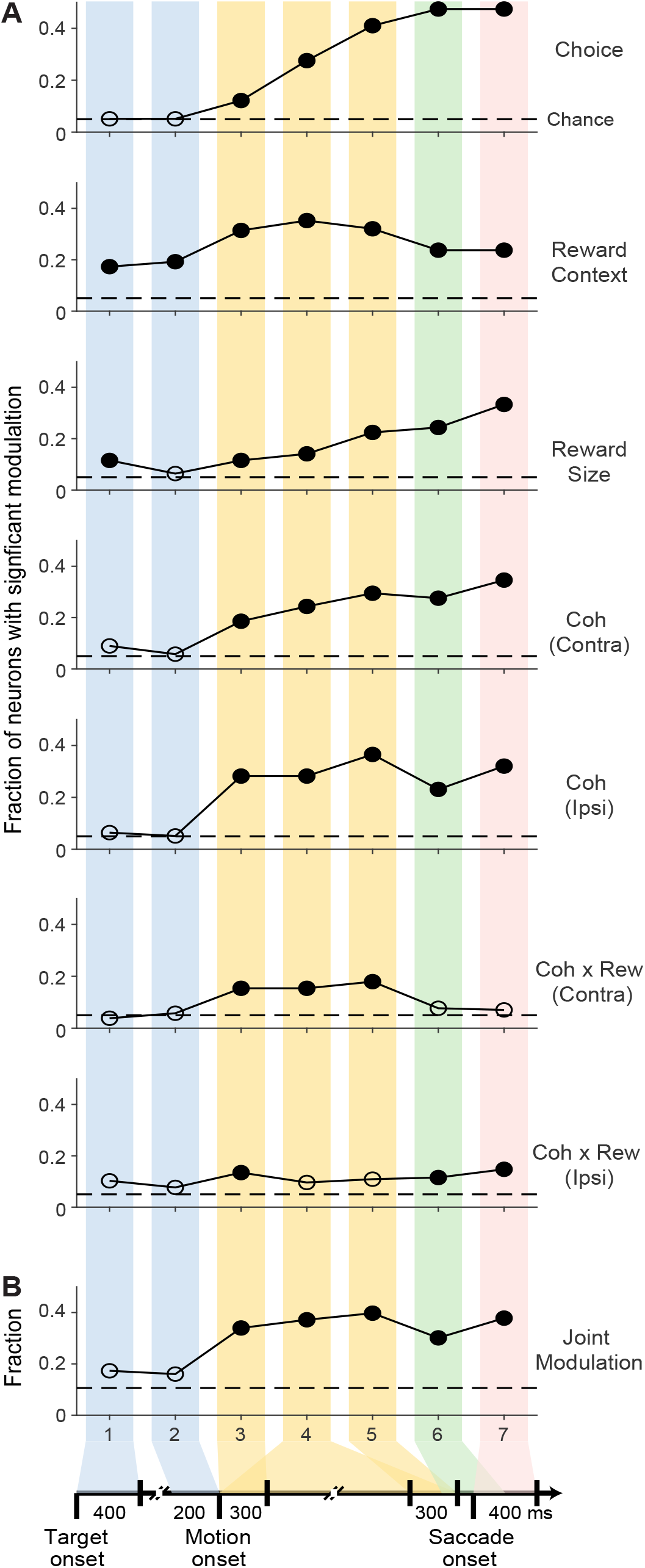
STN activity is modulated by choice, visual evidence, and reward information. A, Fractions of neurons with statistically reliable modulation by task-related factors in the seven task epochs (defined in Figure 1A), as identified by a multiple linear regression (Eq. 2). Horizontal dashed lines: chance levels. Filled circles: fractions that were significantly above chance levels (chi-square test, p<0.05). B, Fractions of neurons with joint modulation, defined as significant modulation by motion coherence (for either choice) and reward context or reward size, as well as significant modulation by the coherence-reward size interaction terms.

Using a previous definition of “joint modulation” (Doi et al., 2020), including modulation separately by motion coherence and reward context or reward size and modulation by the interaction of motion coherence and reward size, we found that ~40% of the neurons showed joint modulation during motion viewing. Moreover, a substantial amount of joint modulation persisted after saccade onset (Figure 3B). These results are consistent with the idea that the STN contributes to the formation and evaluation of complex decisions that incorporate both visual evidence and reward information, at both the single-neuron and population levels.

These modulation patterns of STN neurons for the asymmetric reward task were comparable to those found in the caudate nucleus, another basal ganglia input structure, in the same monkeys performing the same task (Figure 3-S2). These patterns showed more substantial differences with those obtained from a prefrontal region, the frontal eye field (FEF), which showed more prevalent choice modulation, less prevalent reward context modulation during motion viewing, and less prevalent reward size and coherence modulation around/after saccade onset. These inter-regional differences suggest that the basal ganglia are more directly involved than the FEF in mediating decisions that require the incorporation of visual and reward information.

### STN activity encodes multiple decision components

To focus on neurons with the most robust task-relevant activity, we measured firing rates during a baseline period (300 ms before motion onset) and sliding 100 ms windows from motion onset to 150 ms after saccade onset in 50 ms steps. We identified the maximal and minimal z-scores, representing the peak activation and suppression, respectively, for each neuron across all trial conditions (Figure 4C). We applied a threshold of z-score >1.5 for either activation or suppression and focused further analyses on the 87 neurons that met this selection criterion (n = 62 and 25 for monkeys C and F, respectively).

**Figure 4:**
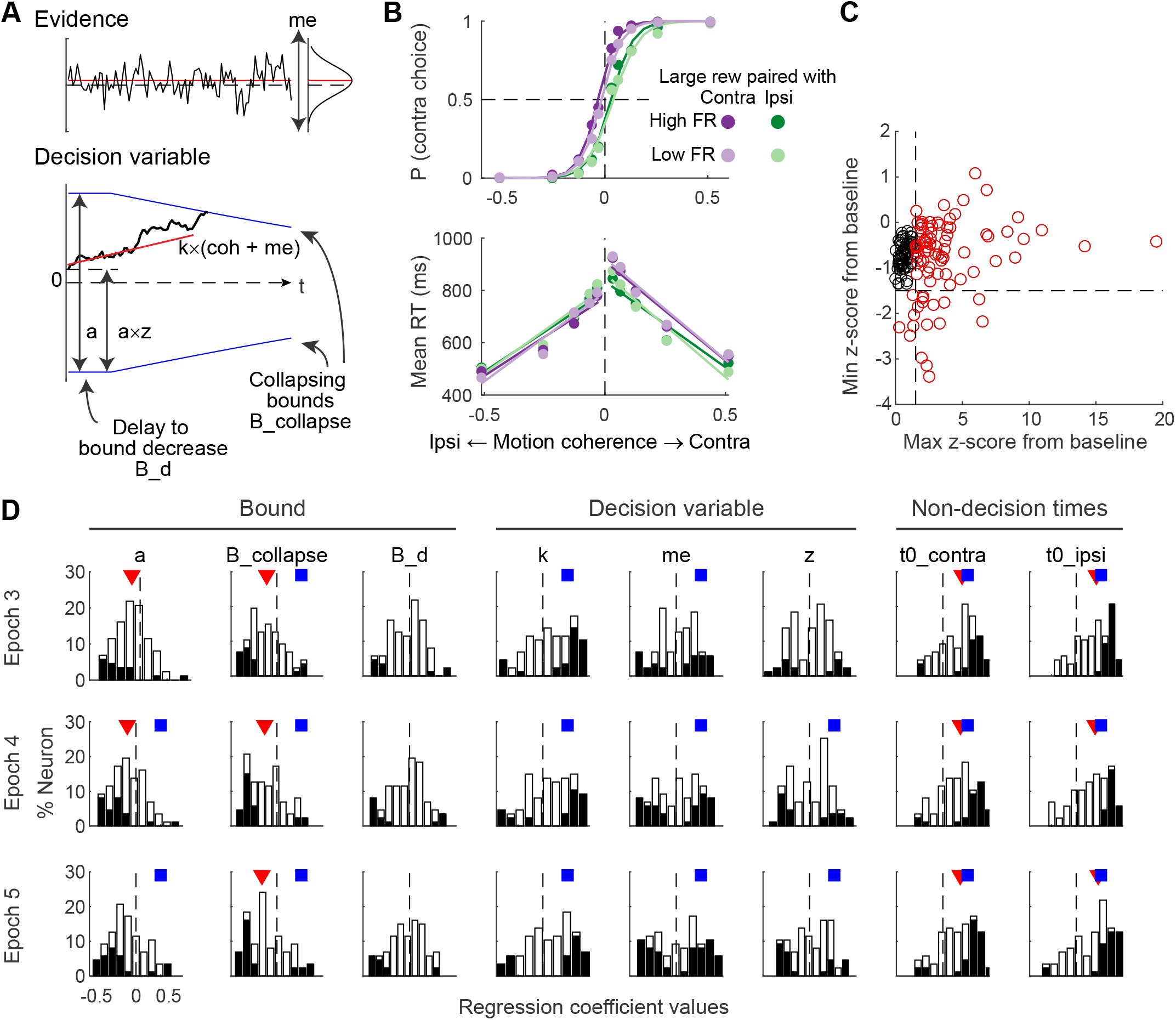
STN activity covaries with multiple DDM parameters. A, Illustration of the DDM. B, Example of average choice and RT performance for trials split by firing rates in Epoch 5. Circles: performance for trials grouped by firing rates and reward contexts (see legend in top panel). Curves: fits by logistic (choice) and linear (RT) functions. C, Identification of neurons with strong task-related modulation. x-axis: z-score of maximal activation for each neuron; y-axis: z-score of maximal suppression for each neuron. Dashed lines: criteria used for identifying task-modulated neurons (modulation z-score>1.5 for activation or suppression). Red circles: neurons included in the analyses in D. D, Histograms of regression coefficients for combinations of activity epochs (rows) and DDM parameters (columns). The parameters include: a, the maximal bound height; B_collapse and B_d, the decay speed and onset specifying the time course of the bound “collapse”; k, a scale factor governing the rate of evidence accumulation; me, an offset specifying a bias in the rate of evidence accumulation; z, an offset specifying a bias in the DV, or equivalently, asymmetric offsets of equal magnitude for the two choice bounds; and t0_Contra and t0_Ipsi, non-decision times for the two choices that capture RT components that do not depend on evidence accumulation (e.g., visual latency and motor delay). Filled bars: neurons showing significant non-zero coefficients (t-test, p<0.05). Red triangles: population median differs from zero (sign rank test), assessed with criteria of p=0.001 (p = 0.05 after multiple comparisons correction for 3 epochs, 8 parameters and both firing rate and firing rate x reward context effects). Blue squares: the proportion of neurons with significant covariation is above chance (chi-square test), assessed with criteria p=0.001.

Many of these STN neurons’ activity covaried with specific computational components of the decision process, quantified as fit parameters of a drift-diffusion model (DDM). The DDM quantifies decision formation in terms of the (possibly biased) accumulation of noisy sensory evidence (the “decision variable”) to a predefined bound that governs choice and RT, which accounts well for behavioral performance on this task (Figure 4A; Fan et al., 2018). As detailed in Methods, parameters *a, B_collapse*, and *B_d* together control the time-dependent bound trajectory, by specifying the initial bound height, the time constant of bound decrease, and the delay to bound decrease, respectively. Parameters *k, me*, and *z* together control the computation of the decision variable, by specifying the scaling factor for evidence, bias in evidence, and bias in the starting value, respectively. Parameters *t0_contra* and *t0_ipsi* specify non-decision times related to additional sensory and/or motor processing for contralateral and ipsilateral choices, respectively.

To probe the behavior-neural relationship, we generated twelve pseudo-sessions for each neuron: for each one of two reward contexts and one of three epochs (epochs 3–5 in Figure 1A, all during motion viewing), we split the trials by the neuron’s median firing rates under those conditions. Figure 4B shows the average choice and RT performance from trials that were split based on neuron’s firing rates in Epoch 5. We fitted the DDM to these pseudo-sessions separately and performed multiple linear regression (Eq. 8) to test for covariations between neural activity (z-scored within each reward context for a given neuron) and each DDM parameter.

We observed several relationships between STN activity and DDM parameters. Higher STN activity during motion viewing was associated with lower initial bound height and slower bound decreases (Figure 4D; *a* and *B_collapse*). Higher STN activity was also associated with longer non-decision times for both choices (*t0_contra* and *t0_ipsi*). Although the activity of a substantial proportion of STN neurons covaried with parameters associated with the decision variable (*k, me*, and *z*), there was no consistent directionality (filled bars were present for both positive and negative regression coefficients). There was no consistent relationship between DDM parameters and activity-reward context interaction across the population, but for a substantial proportion of neurons, there were reliable covariations in either direction for parameter *me* (Figure 4-S1). These results suggest that STN neurons as a population may be involved in modulating multiple computational components in the DDM framework. We next examined if and how these modulations related to the different activity patterns evident in Figures 2 and 3.

### STN subpopulations encode different components of decision formation

We identified three clusters of STN neurons (including only those neurons that met the threshold for task-relevant activation) using k-means and linkage-based clustering analyses. We represented each neuron with a 328-dimensional vector summarizing its normalized firing rates in 14 task periods for 20 trial types (two choices, two reward contexts, and five coherence levels), as well as the regression coefficients for the split-trial DDM fitted values above (eight values each for the main firing rate effects and the firing rate-reward context interaction effects, in three epochs). We trimmed 10–12 neurons that were “outliers” that did not belong to major subpopulations (see Methods). Inspection of the dendrogram (hierarchical cluster tree) suggested that our STN samples can be reasonably grouped into three clusters, although other groupings are possible using different clustering cutoffs (Figure 5-S1).

The clustering results appeared stable and robust. The clustering results from k-means (Figure 5) and linkage-based (Figure 5-S2) methods were highly similar, with a Rand index of 0.93. Clustering of data from both monkeys combined compared to each monkey considered separately had mean rand index values of 0.94 and 1 for monkeys C and F, respectively (i.e., neurons from one monkey tended to be assigned to the same cluster regardless of whether the clustering was based on data from that monkey alone or both monkeys together), indicating comparable cluster boundaries for the two monkeys. The average firing rate patterns of clusters identified from each monkey separately are shown in Figure 5-S3 for comparison. The quality of clustering, as assessed with silhouette score and Rand index, was robust and stable even when only a subset of trials was used to compute firing rates for each neuron (Figure 5-S4). These clusters were also relatively stable between task types: assuming that there are three clusters, the rand index was 0.92 and 0.86 between clustering results based on neural activity on equal- and asymmetric-reward motion discrimination tasks, using the k-means and linkage methods, respectively. The clustering results were primarily driven by activity patterns: clustering based on the firing rates only (a 280-D vector per neuron) reproduced essentially the same results (the Rand index between the two sets of clusters was 1 and 0.92, for the k-means and linkage methods, respectively).

**Figure 5:**
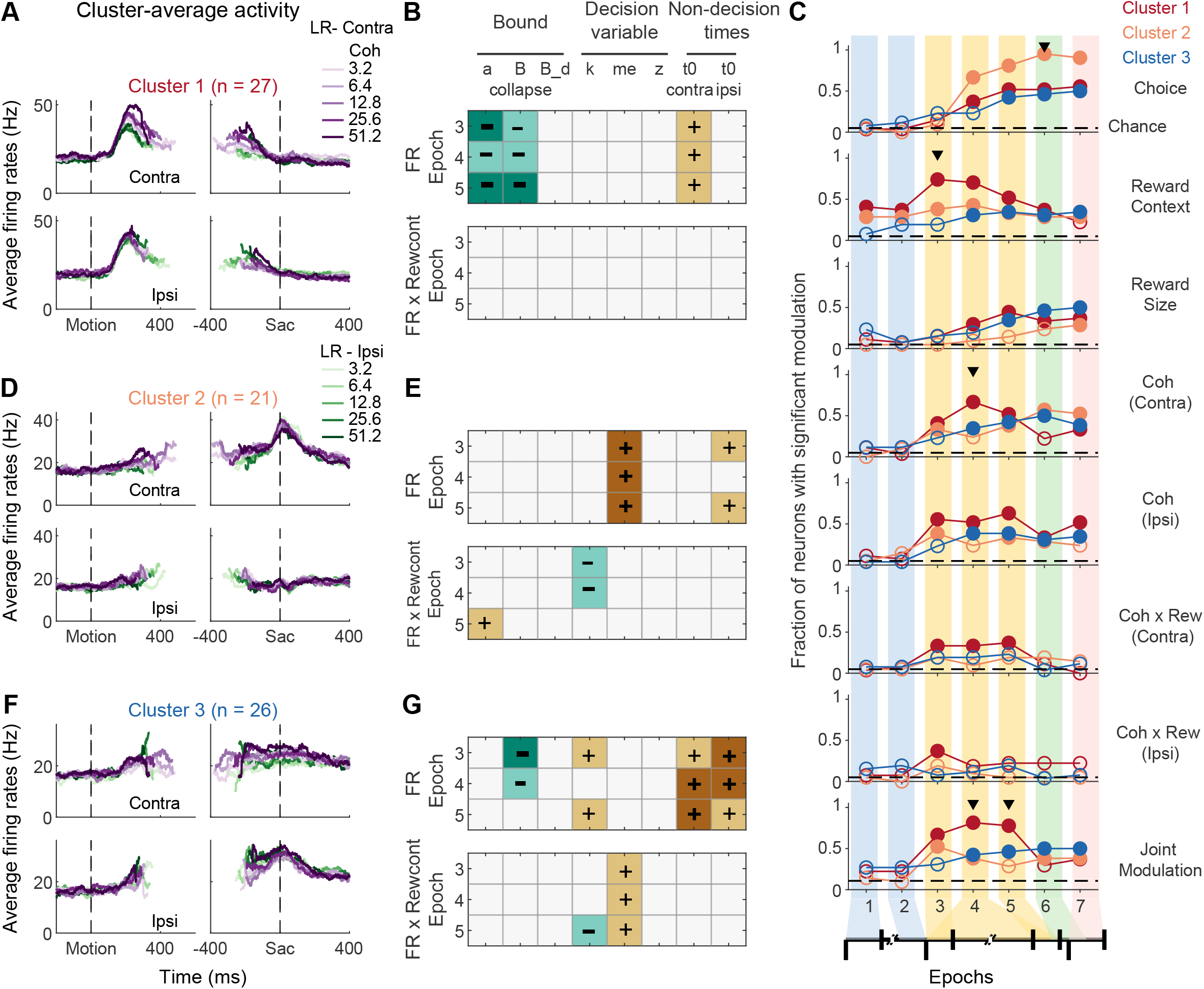
STN consists of subpopulations with distinct activity patterns and relationships with DDM components. A,D,F, Average firing rates for each cluster of neurons for asymmetric-reward trials, separately for correct trials with contralateral (top) and ipsilateral (bottom) choices and aligned to motion (left) and saccade (right) onsets. Colors indicate coherence levels and reward context (see legend). B,E,G, Relationship between activity measured in the three epochs and DDM components for each cluster. Top: sign-test results for the null hypothesis that the regression coefficients for the effect of firing rate in the three epochs on the DDM component have a zero median. Bottom: sign-test results for the null hypothesis that the regression coefficients for the reward context-firing rate interaction have a zero median. Dark brown/teal: positive/negative median with p<0.001. Light brown/teal: positive/negative median with p<0.05. C, Fractions of neurons with significant modulation by task-related factors in the seven task epochs, separately for each cluster. Same format as Figure 3. Horizontal dashed lines: chance levels. Filled circles: fractions that were significantly above chance levels (Chi-square test, p<0.05). Triangles: epochs in which there was a significant difference among clusters (Chi-square test, p<0.05/7 epochs).

The three clusters of neurons differed in their task-related activity patterns. The first cluster of neurons showed early activation that gradually returned to baseline during motion viewing (Figure 5A). These neurons were more likely to be sensitive to reward context during motion viewing (Figure 5C, “reward context” panel). The second cluster showed consistent choice selectivity for the contralateral choice (Figure 5D), especially around saccade onset (Figure 5C, “choice” panel). The third cluster showed late-onset activation, with moderate choice modulation also emerging during late motion viewing (Figure 5F). When the proportion of neurons reached above chance level for any decision-related modulation, the timing for this cluster tended to lag behind the other two clusters (Figure 5C).

These clusters also had distinct relationships to the fit DDM parameters, based on sorting the results from the pseudo-session analyses described above. Cluster 1 activity was most consistently associated with reward context-independent bound adjustments, with higher activity corresponding to lower initial bound heights (*a*) and slower decreases in bounds (*B_collapse*; Figure 5B, teal-colored boxes), along with a weaker relationship with non-decision times for contralateral choices (light brown boxes). Cluster 2 activity was most consistently associated with a larger bias in evidence (*me*) toward the contralateral choice (Figure 5E, top panel), along with weaker relationships with the ipsilateral non-decision time (*T0_Ipsi*), the context-dependent decision variable scale factor (*k*), and the context-dependent changes in bound (Figure 5E, bottom panel). Cluster 3 activity was most consistently associated with slower decreases in bounds and non-decision times for both choices, along with other mostly weaker relationships to various other decision-related parameters that in many cases depended on reward context (Figure 5G). Histograms corresponding to the teal and brown boxes are shown in Figure 5-S5.

Because the pseudo-session analysis normalized firing rates within each reward context, *bRew* in Eq. 8 can be used to detect whether both neural activity and behavior are sensitive to reward context, but cannot test whether stronger reward context-modulation in neural activity is associated with larger behavioral changes. We therefore performed a separate correlation analysis to test for a relationship between reward context modulations of neural activity and DDM parameters, focusing on possibilities corresponding to the dark brown or teal boxes in Figure 5. No reliable relationships were found for Clusters 1 and 3. In contrast, Cluster 2 neurons exhibited positive relationships between reward context modulation of activity and reward context modulation of *me* values (Figure 6 and 6-S1). This result suggests that this subpopulation may contribute to the implementation of reward-dependent choice biases that involve how the evidence is used to form the decision.

**Figure 6:**
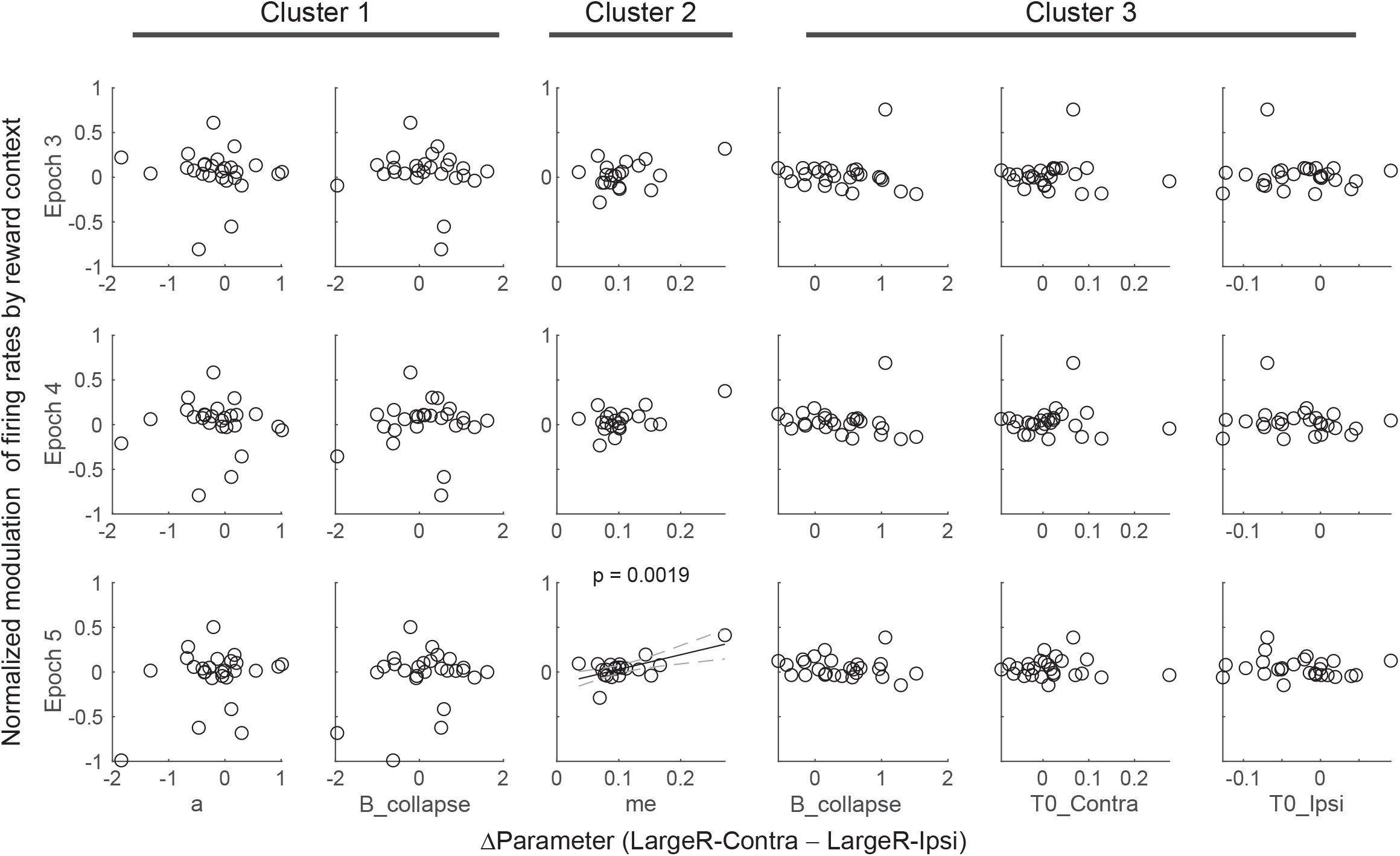
Relationship between reward context modulation of firing rates and reward context modulation of DDM parameters, based on k-means clustering. Each panel shows the scatterplot of reward context modulation of firing rates in an epoch, normalized by the average firing rate in the same epoch, and the difference in a DDM parameter value between the two reward contexts. Each data point represents one neuron in a given cluster. Significant correlation (t-test, p<0.05/18) is indicated by the raw p value and regression line with confidence intervals.

Thus, the STN’s diverse functions may involve distinct contributions to both decision formation and non-decision (e.g., basic visual and/or motor) processes by functionally defined subpopulations.

### STN subpopulations encode signals related to decision evaluation

We used the asymmetric-reward manipulation to partially decorrelate two signals that can be used to evaluate decisions, namely choice accuracy and reward expectation (Fan et al., 2024). Choice accuracy describes the probability that a choice is correct given the evidence. Reward expectation describes the expected reward given a choice. Following our previous approaches, we computed the partial Spearman correlation between average activity in a given time window and each of the two quantities (choice accuracy and reward expectation), while controlling for the other.

Choice accuracy and/or reward expectation were reflected in the peri-saccadic activity of many of the STN neurons in our sample (Figure 7A). Overall, STN neurons whose activity showed significant correlation with choice accuracy or reward expectation were similarly prevalent (Figure 7C, upper panel). More neurons tended to show activity that was positively correlated with reward expectation than with choice accuracy, especially after saccade onset (Figure 7C, lower panel). This trend was significant when assessed at *p*<0.05 but did not survive multiple comparison correction. All three subpopulations included neurons with evaluative signal representation (Figure 7B, D), with similar prevalence. The tendency of positive correlation for reward expectation seemed to be driven mostly by the first subpopulation. However, the small sample sizes did not offer enough power to properly assess potential subpopulation differences. Thus, both choice accuracy and reward expectation signals were present in STN activity, with potential differences in how they are represented among the functionally defined subpopulations.

**Figure 7:**
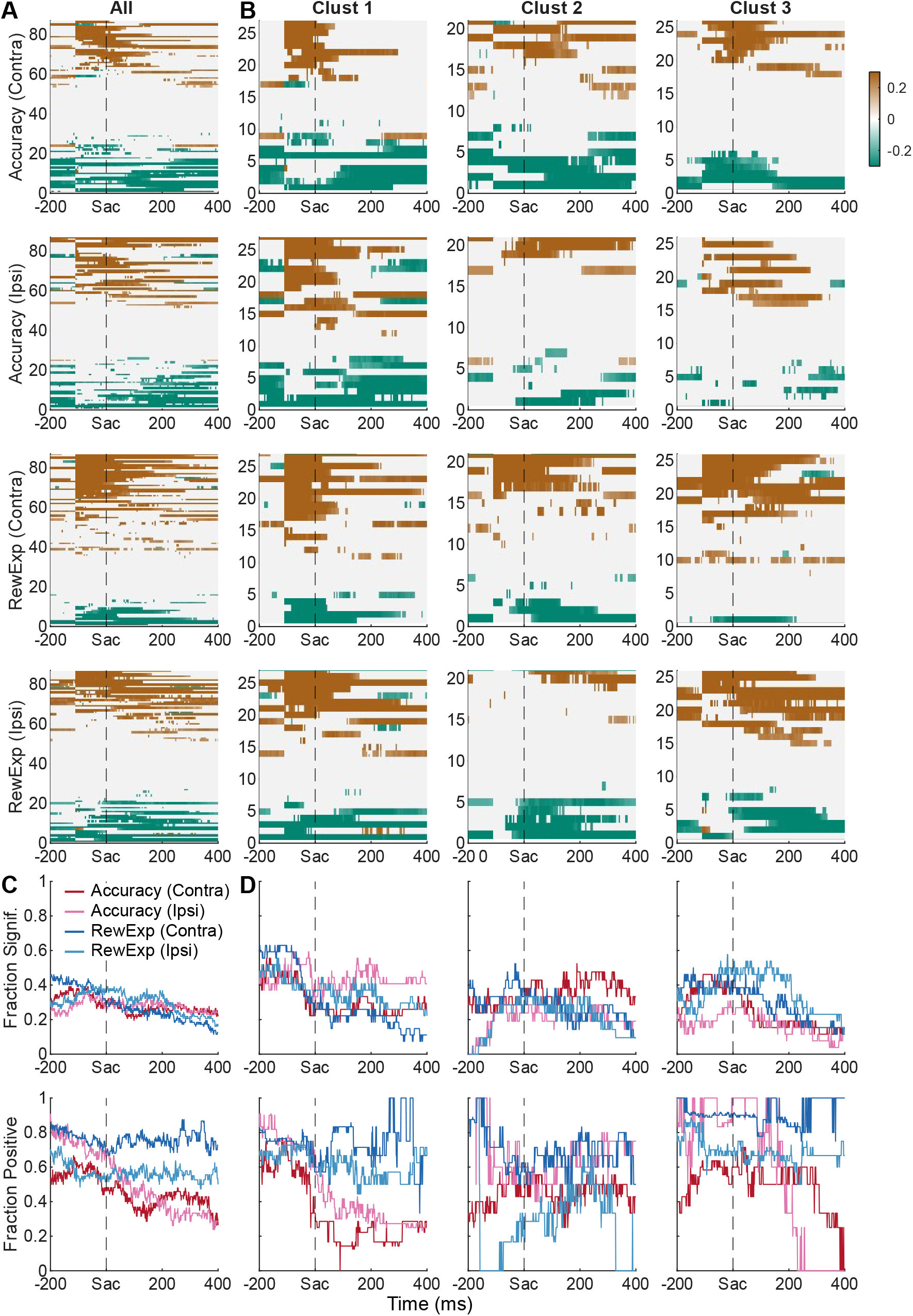
Activity of STN neurons reflect decision evaluation signals. A, Heatmaps of partial correlation coefficients between neural activity and post-decision choice accuracy and reward expectation for contralateral and ipsilateral choices, after accounting for the effects of reward expectation and choice accuracy, respectively. Values that did not differ from 0 (t-test, p>0.05 after multiple comparison corrections) were set to zero. Only neurons with task-modulated activity were included (*n*=87). B, Heatmaps for each cluster (column). Clusters were defined using the k-means method (same as Figure 5). C, Fractions of neurons with non-zero partial correlation (top) and with positive partial correlation (bottom). D, Fraction plots for each cluster.

### STN subpopulations are intermingled

In our previous study of STN activity using the equal-reward motion discrimination task, we did not find any systematic anatomical organization of the different STN subpopulations. Here we found a similar anatomical intermingling of neurons from the three clusters when tested using the asymmetric-reward version of the task (Figure 8). In particular, the mean Silhouette scores (using Euclidean distance) between neurons with robust task modulation and those without were −0.011 and 0.07 for the two monkeys, respectively, indicating strong overlap between the two groups (Figure 8A). Similarly, the mean Silhouette scores among the clusters were −0.02 and 0.097 for the two monkeys, respectively, indicating that the different subpopulations were intermingled (Figure 8B).

**Figure 8:**
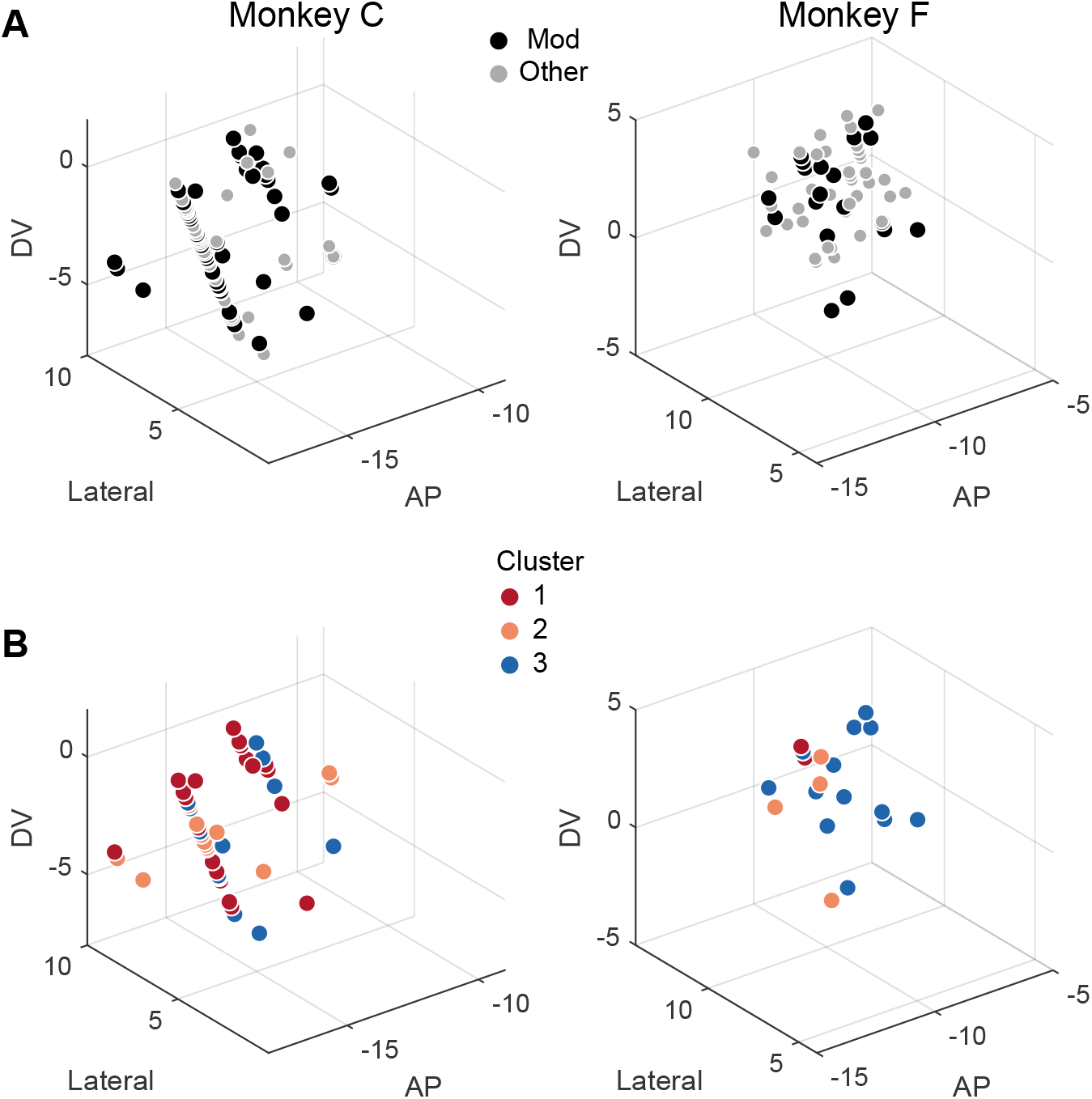
STN neurons with different activity patterns are intermingled. A, Locations of neurons in the two monkeys. Black points are neurons considered “task-modulated”. B, Locations of neurons in different clusters. Distance values are relative to the anterior commissure. AP: anterior/posterior; DV: dorsal/ventral.

## Discussion

We used an asymmetric-reward random-dot visual motion discrimination task to better understand the role of the monkey STN in the formation and evaluation of complex decisions that rely on both sensory and reward information. We found that the STN is functionally heterogeneous, with neurons that exhibit a diversity of task-related activation patterns and relationships to behavior. This functional diversity, along with a lack of clear anatomical organization, is consistent with the multiple effects of STN stimulation in patient populations on decision-making and our previous results in monkeys, including reductions in response times, a weaker dependence on evidence, and changes in the maximal value and trajectories of the decision bound (Frank et al., 2007; Cavanagh et al., 2011; Coulthard et al., 2012; Green et al., 2013; Zavala et al., 2014; Herz et al., 2016; Pote et al., 2016; Branam et al., 2024). Building on our prior findings from the same monkeys performing an equal-reward version of the task, we identified a functional organization of this diversity, including three clusters of STN neurons with distinct putative roles in the decision process and potentially different connectivity patterns (Figure 9), as follows.

**Figure 9:**
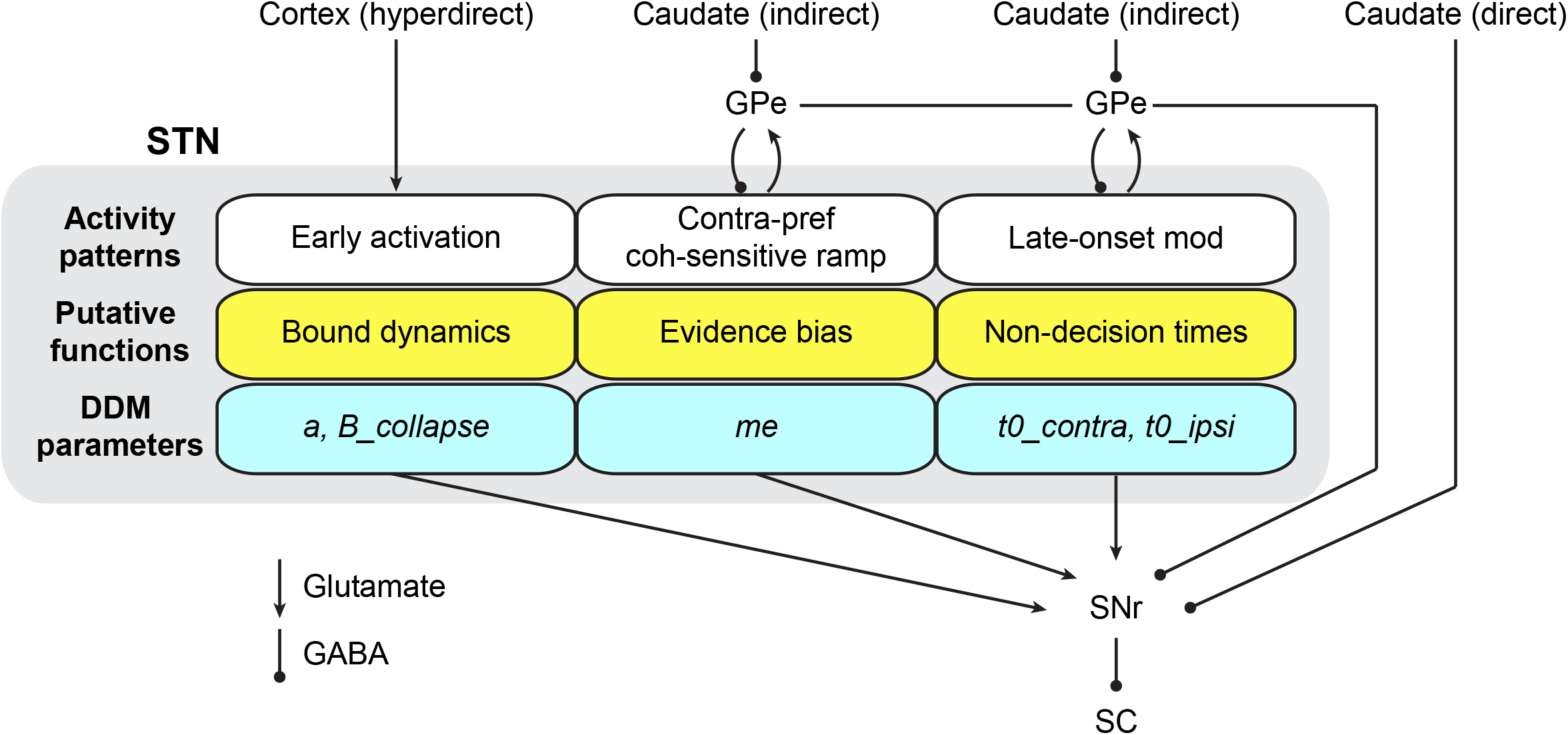
Hypothesized functions and connectivity of STN subpopulations. The three STN subpopulations may receive different inputs and serve distinct functions. Cluster 1 neurons may receive direct input from the cortex via the hyperdirect pathway and provide early-onset modulation of bound dynamics. Cluster 2 and 3 neurons may receive inputs from different GPe subpopulations in the indirect pathway, which relay caudate signals related to evidence accumulation and saccade generation, to mediate evidence bias and non-decision times, respectively. These different STN outputs and the outputs from GPe and the caudate direct pathway converge onto SNr to affect decision-related SC activity via inhibition. Abbreviations: GPe: external segment of globus pallidus; SNr: substantia nigra pars reticulata; SC: superior colliculus.

The first group (Cluster 1 here, 2 in the prior study) included neurons that tended to be most active early in the decision process then return to around baseline levels by the time of the saccadic response. Activity of these neurons was most closely associated with session-by-session differences in the decision bound, including its overall height and rate of collapse over time (parameters *a* and *B*_*collapse*_ from the DDM fits). This role in setting the decision bound contrasts with our previous findings from the caudate, which is a source of the indirect pathway that provides one set of inputs to the STN, where we found neurons with strong choice selectivity but no clear representation of a decision bound (Ding and Gold, 2010). We therefore speculate that neurons in this group may receive input from cortex via the hyperdirect pathway and not via the striatum (including the caudate) to control reward context-independent adjustments to the decision bound.

The second group (Cluster 2 here, 1 in the previous study) included neurons with the strongest choice selectivity. Activity of these neurons was most closely associated with differences in biases, in particular a reward-dependent scaling of the evidence that in the DDM gave rise to faster and more prevalent choices to the high-reward alternative (parameter *me*). Because we also found that the activity of neurons in caudate can encode and affect reward-dependent biases (Doi et al., 2020), we speculate that neurons in this group may also receive input via the indirect pathway that originates in the caudate to contribute to certain context-dependent adjustments of the decision variable.

The third group included neurons that tended to be most active late in the decision process, with moderate selectivity for reward, coherence, and choice emerging around the time of choice. Activity of these neurons was most closely associated with differences in non-decision times, which can include extra sensory and/or motor processing beyond what is accounted for by the DDM (parameters *t0*_*contra*_ and *t0*_*ipsi*_ from the DDM fits). We also found substantial effects of STN microstimulation on non-decision times in our previous study, which was likely mediated in large part via neurons in this group (Branam et al., 2024). Because we also found that perturbation of caudate activity can affect non-decision times (Ding and Gold, 2012a), we speculate that neurons in the indirect pathway contribute to visual/motor processing independent of decision formation and that this group of STN neurons may receive input via the indirect pathway that originates in the caudate.

These findings also provide new insights into how STN’s contributions to decision formation relate to those used in various computational models. As described in our previous study (Branam et al., 2024), the neural activity patterns corresponding to the three clusters are reminiscent of predictions of three separate models: (Ratcliff and Frank, 2012) for the current Cluster 1, (Bogacz and Gurney, 2007) for Cluster 2, and (Wei et al., 2015) for Cluster 3. We used microstimulation in that study to probe relationships between STN activity and specific decision-making functions. However, we found few systematic relationships between DDM-derived functions and cluster identity of the neuron recorded at the site of microsimulation, likely because microstimulation affects an extended volume of brain tissue that includes intermixed neurons from different clusters. Here, we took a different approach and related DDM parameters to modulations of single-unit responses. We found that, as we previously inferred, Cluster 1 neurons appear to play roles similar to that found in the model by Ratcliff and Frank (2012), involving bound adjustments. However, functions of the other two clusters do not seem to map as neatly onto those found in the other two models. Furthermore, our clustering analysis aimed to identify common activity profiles in the STN population, while leaving behind many neurons that either did not show consistent task-related modulation or had less common activity profiles (e.g., those that were far from others in the vector space and those with too infrequent occurrence to form detectable clusters). More work is needed to continue to refine our understanding of the specific computational contributions of the STN to decision formation.

These contributions may involve adjusting decision biases like those resulting from asymmetric rewards in our study. A substantial fraction of STN neurons in all three subpopulations were sensitive to reward context. This sensitivity including relatively weak relationships between interactions between reward context and neural activity and the scaling factor (*k*) and evidence bias (*me*) for Clusters 2 and 3 neurons (Figure 5E and G, bottom panels) and relationships between the magnitude of reward-context modulation and *me* bias for Cluster 2 neurons (Figure 6). Although Cluster 1 neurons had the highest prevalence of reward-context modulation, we did not observe any behavioral correlates of such modulation (Figure 5E, bottom panel; Figure 6). The exact contributions of these subpopulations are challenging to elucidate, as their intermingled localization make common perturbation techniques, such as electrical microstimulation or optogenetic manipulations, not suitable. It would be interesting to examine if these subpopulations differ in molecular or connectivity properties (e.g., as we speculated above) that can be utilized to precisely target each subpopulation.

The prevalence of joint modulation by reward and evidence-related factors in our sample was comparable to what we observed in the caudate (Figure 3-S2; Doi et al., 2020; Fan et al., 2020), suggesting that the reward-biased decision behavior reflects interactions among multiple BG nuclei (and likely other brain regions). Both the caudate nucleus and STN receive dopaminergic innervation from the substantia nigra pars compacta (Galvan et al., 2014), which may carry reward-related signals, and cortical projections, which may carry evidence accumulation-related signals (Bogacz and Gurney, 2007; Ding and Gold, 2012b; Ding, 2015; Wei et al., 2015). These inputs may be combined in the STN to support the generation of a common-currency decision variable. Alternatively, it is possible that the STN receives already combined signals either directly from the cortex or indirectly from the caudate projection to the external segment of globus pallidus, which has strong reciprocal connections to the STN. Future work is needed to understand how different BG nuclei obtain decision-related signals that reflect incorporation of reward information and noisy evidence accumulation. Furthermore, both caudate and STN outputs are sent to the output BG nucleus, specifically the SNr, for oculomotor decisions. It remains an open question how the various forms of decision-related signals from multiple BG sources are combined in the BG output to affect the eventual decisions.

Our sample was selected based on neural activity on the visual motion discrimination task. The high fraction of neurons showing joint modulation by reward and noisy sensory evidence in this sample suggested that these STN neurons may participate in a variety of decisions that involve either or both of these factors. It remains to be tested whether the same STN neurons are also involved in decisions or movement modulation based on reward properties alone (Zaghloul et al., 2012, 2012; Espinosa-Parrilla et al., 2015; Zénon et al., 2016; Nougaret et al., 2022). The early-onset activity profile of Cluster 1 neurons is particularly intriguing for its possible correspondence to previously identified STN neurons that may mediate response inhibition in stop-signal (countermanding) tasks (Schmidt et al., 2013; Pasquereau and Turner, 2017) or STN neurons that may mediate switching from automated to controlled actions (Isoda and Hikosaka, 2008).

In summary, we characterized single-neuron activity in the STN of monkeys performing a decision task that requires incorporation of noisy visual evidence and different reward context information. These results advanced our understanding of the roles of the STN in decision making, by refining previous hypotheses about its role in the evidence-accumulation-to-bound framework and, for the first time, revealing how activity of different STN subpopulations were modulated to contribute to the formation and evaluation of reward-biased decisions.

## Methods

We used two adult male rhesus monkeys (*Macaca mulatta*) that were trained extensively on the asymmetric-reward visual motion direction-discrimination (dots) task. All training, surgery, and experimental procedures were in accordance with the National Institutes of Health Guide for the Care and Use of Laboratory Animals and were approved by the University of Pennsylvania Institutional Animal Care and Use Committee (protocol # 804726).

### Task design

The behavioral task (Figure 1A) has been described in detail previously (Fan, 2018). Briefly, we trained the monkey to report the perceived motion direction of the random-dot stimulus with a saccade at a self-determined time. The monkey’s choice of saccade was rewarded if it was congruent with the motion direction. The monkey’s eye position was monitored with a video-based eye tracker and feedback for a fixation break or choice error was given based on online comparisons between the monkey’s eye position and task-relevant locations. Motion directions and five levels of motion coherence (defined as the fraction of dots moving coherently in the same direction) were randomized across trials. For the equal-reward dots task, all correct saccades were rewarded with the same medium amount of juice. For the asymmetric-reward dots task, the saccade-reward associations (reward contexts) were alternated in blocks of trials, such that in a given block, a correct saccade to one choice target was paired with a large amount of juice and a correct saccade to the other target was paired with a small amount of juice. Block changes were signaled to the monkey with color cues.

### Electrophysiology

The general surgical and data-acquisition methods were described in detail previously (Ding and Gold, 2010, 2012a). Neural activity was recorded using glass-coated tungsten electrodes (Alpha-Omega) or polyamide-coated tungsten electrodes (FHC, Inc.), driven by a NAN microdrive (NAN Instruments, LTD) mounted on a grid system. STN was localized using the same criteria as described previously (Branam 2024), by comparing MRI scan images, recording coordinates, and nearby landmark locations (thalamus, reticular nucleus of the thalamus, zone incerta, and SNr) and by assessing neurons’ baseline firing patterns.

### Initial data processing and screening

Saccade reaction time (RT) was measured offline with established velocity and acceleration criteria. Trials were excluded if the online and offline detection of choice-indicating saccades were mismatched (e.g., the saccade did not follow the stereotypical trajectory). Single neurons were identified by offline spike sorting (Offline Sorter, Plexon, Inc.). Neurons with fewer than five correct trials per choice × coherence × reward context combination were excluded.

### Behavioral analysis

To quantify reward context-induced biases, a logistic function was fitted to the choice data for all trials for each session:

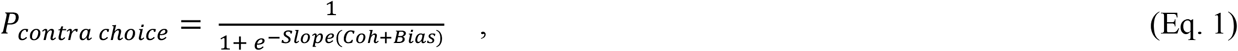

where *Coh* is the signed motion coherence,

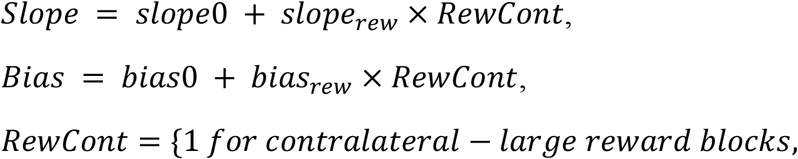

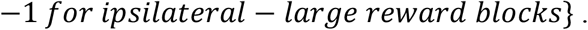

To quantify decision-related computations that account for both choice and RT, we fitted a drift-diffusion model (DDM; Figure 5A) to these data simultaneously, following previously established procedures (Fan et al., 2018). Briefly, the DDM assumes that motion evidence is accumulated over time into a decision variable (DV), which is compared to two choice bounds that decrease in magnitude (“collapse”) over time within a trial. A choice is made when the DV crosses either bound, such that the time of crossing determines the decision time and the identity of the bound determines the choice identity. The model has eight basic parameters, including: 1) *a*, the maximal bound height; 2) *B_collapse* and *B_d*, the decay speed and onset specifying the time course of the bound “collapse”; 3) *k*, a scale factor governing the rate of evidence accumulation; 4) *me*, an offset specifying a bias in the rate of evidence accumulation; 5) *z*, an offset specifying a bias in the DV, or equivalently, asymmetric offsets of equal magnitude for the two choice bounds; and 6) *t0_Contra* and *t0_Ipsi*, non-decision times for the two choices that capture RT components that do not depend on evidence accumulation (e.g., visual latency and motor delay). DDM model fitting was performed separately for each session, using the maximum *a posteriori* estimate method (python v3.5.1, pymc 2.3.6) and prior distributions suitable for human and monkey subjects (Wiecki et al., 2013). We performed at least five runs for each variant and used the run with the highest likelihood for further analyses.

To quantify the expected choice accuracy and reward expectation, we used previous methods (Fan, 2024). Specifically, we defined choice accuracy as the estimated accuracy, on average, given the current choice and decision time (DT), as following:

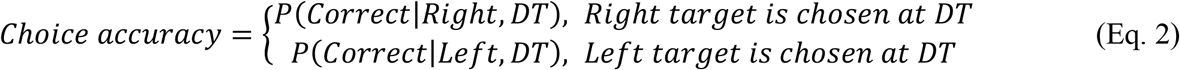

where DT is the decision time that equals RT minus non-decision time (estimated from DDM fits). The righthand side was computed by marginalizing over all possible coherences. For example, for Right choices,

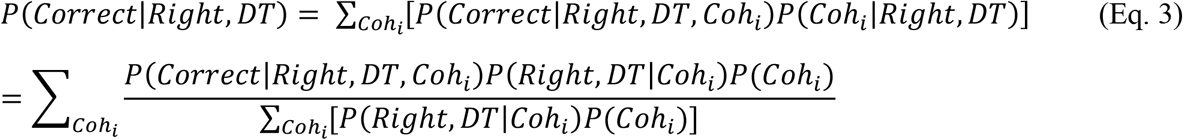

By task design,

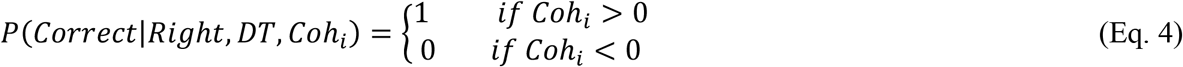

*P* (*Right, DT* | *Coh*_*i*_) was obtained by numerical simulation of the DDM using the best-fitting parameters. For each coherence, we obtained the probability of the decision variable (DV) attaining a value *x* at time *t,pdf* _*DV*_ (*t*) = *P*(*DV* (*t*) = *x* | *Coh*_*i*_), using the best fitting DDM parameters of each session and reward context.

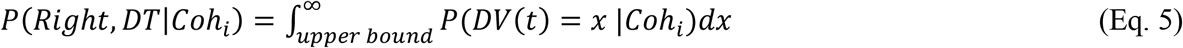

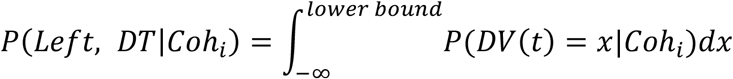

After obtaining an estimate of choice accuracy,

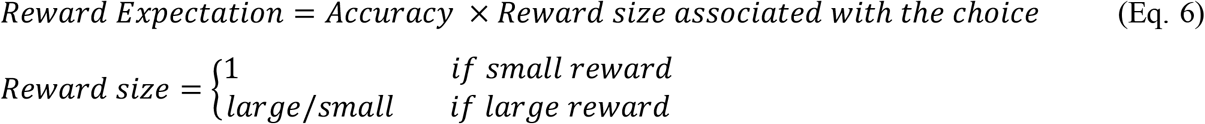

### Neural activity analysis

All analyses were performed on activity on the asymmetric-reward task. Only correct trials were included for neural activity analysis to avoid the confound of potential task disengagement during error trials. To relate activity to decision-related factors, we performed two regression analyses. First, for each single unit, we computed the average firing rates in seven task epochs (Figure 1A): 1) a 400 ms window beginning at target onset (focusing on visual response to target presentation), 2) a 200 ms window ending at dots onset (focusing on activity before motion viewing), 3) a 300 ms window beginning at 100 ms after dots onset (focusing on early motion viewing), 4) a variable-duration window from 100 ms after motion onset to 100 ms before saccade onset (focusing on the whole motion viewing period), 5) a 300 ms window ending at saccade onset (late motion viewing, pre-saccade activity), 6) a 300 ms window beginning at 100 ms prior to saccade onset (peri-saccade activity), and 7) a 400 ms window beginning at saccade onset (post-saccade activity). We performed a multiple linear regression on the spike counts from correct trials, for each task epoch separately.

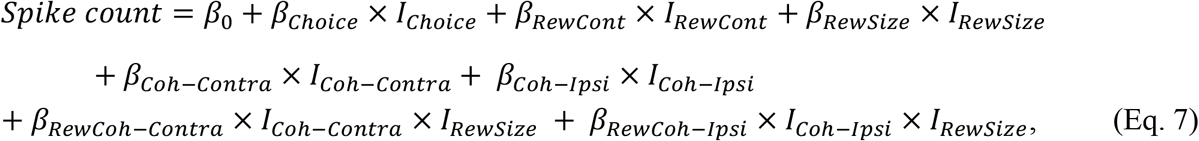

where

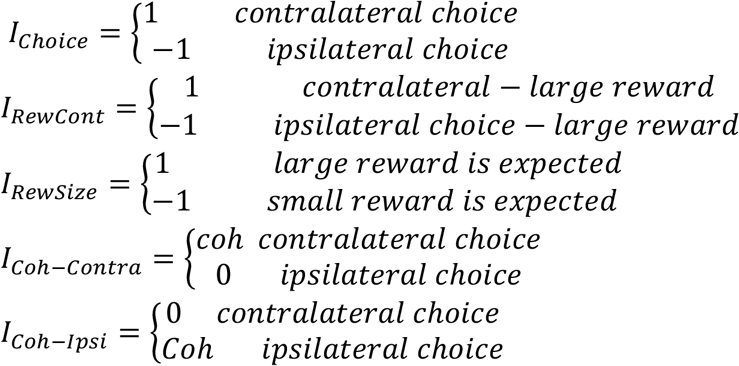

Significance of non-zero coefficients was assessed using *t*-test (criterion: *p* = 0.05).

Second, for each single unit, we also performed running regressions using Eq. 7 on the spike counts within 150 ms windows with 10-ms steps. These running regressions were performed on activity aligned to target, motion, and saccade onsets separately. Only correct trials were included. Time windows with <10 correct trials were excluded.

### Relate neural activity to DDM components

For each neuron and reward context, we split the trials by the median firing rates in one of the three epochs that covered the motion viewing period (i.e., Epochs 3–5). We fitted the DDM separately to trials with different high/low firing rates × reward context combinations, resulting in 12 sets of fitted values. We then measured the average firing rates in the three epochs and z-scored these values separately for each reward context × epoch combination. We used this z-scoring approach to control for the potential confound of observing activity-behavior covariation simply because both are modulated by reward context. We used a linear regression to assess the influences of reward context, firing rates, and their interactions on each DDM parameter using the regression:

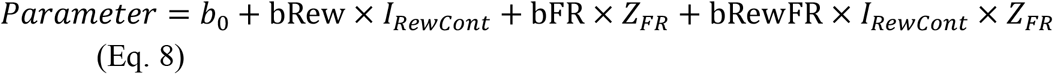

T-test was performed to test whether a regression coefficient differed from zero for each neuron (criterion: *p*=0.05). Sign test was performed to test whether median values of regression coefficients differed from zero for the population (criterion: p = 0.05 and 0.001 for without and with multiple comparison corrections, respectively). Chi-square test was performed to test whether a significant portion of the population showed significant non-zero coefficients, regardless of sign (criterion: p = 0.05 and 0.001 for without and with multiple comparison corrections, respectively).

### Cluster analysis

We represented each neuron with a vector containing: 1) 280 data points measuring the average firing rates within fourteen 100-ms windows from 300 ms before motion onset to 150 ms after saccade onset (in 50 ms steps) for twenty trial conditions (i.e., combinations of two choices, five coherence levels, and two reward contexts), and 2) 48 data points measuring the regression coefficients from Eq. 8 for the split-trial analysis above (16 coefficients for each of the three epochs). The firing rates were z-scored based on pre-motion baseline firing rates for each neuron across all trial conditions.

We used two methods for clustering these vectors using the correlation distance metric (1-correlation between vectors). The first method was the linkage method based on agglomerative hierarchical cluster trees. To identify outlier neurons that did not belong to major subpopulations, we performed a linkage clustering using 0.7 as the initial cutoff (Figure 5-S1A) and trimmed neurons that were grouped into clusters with fewer than three members. Such trimming filtered out 10 neurons (8 and 2 from monkeys C and F, respectively). We then re-computed the tree and performed clustering using 0.85 as the final cutoff (Figure 5-S1B), resulting in three clusters.

The second method was the k-means clustering method. To identify outlier neurons, we computed pair-wise correlations between neurons and trimmed neurons with a maximal correlation value of <0.5 (i.e., these neurons were not similar to any other neurons). Such trimming filtered out 12 neurons (8 and 4 from monkeys C and F, respectively). We then performed k-means clustering on the remaining neurons, assuming three clusters.

To quantify the consistency between two runs of clustering, we computed the Rand index as the number of neuron pairs with consistent grouping (i.e., they were placed in the same cluster for both runs or they were placed in different clusters for both runs), normalized by the total number of possible neuron pairs. A value of 1 indicates that the two clustering runs produce identical results, and a value of 0 indicates that the two runs do not agree on any pairs of neurons.

To quantify the separation of clusters, we computed silhouette scores as the difference between mean intra-cluster distance and the mean nearest-cluster distance, normalized by the maximum of the two values. A positive score indicates that the member is closer to its same-cluster neighbors than different-cluster neighbors. Clustering runs with high mean silhouette score were considered to have better cluster separation.

We assessed the robustness of the clustering results in two ways. First, clustering was performed on both monkeys’ data and on each monkey’s data separately (Figure 5-S3). For each monkey’s neurons, we measured the Rand index between cluster identities obtained from clustering just the monkey’s data and those obtained from clustering both monkeys’ data. High Rand index values indicate that similar cluster boundaries are present for both monkeys and the clusters do not simply reflect idiosyncratic properties of a monkey’s STN population. Second, clustering was performed based on firing rates vectors measured from a subset of trials. For each iteration and fraction of trials, we randomly resampled correct trials with replacement for each neuron. Because the monkey made fewer correct choices for low-coherence, small-reward trials, we repeated resampling until at least one trial was included for every trial type (i.e., a vector with the same 280 dimensions can be generated). Ten iterations of resampling were performed for each fraction-of-trials value, the resampled vectors were clustered into three clusters, and the mean Silhouette scores and between-iteration Rand index were computed as above (Figure 5-S4).

### Neural activity analysis: relating activity to decision-evaluation signals

To relate a neuron’s activity to decision-evaluation signals, for each trial, we computed the choice accuracy and reward expectation and measured the firing rates in running 300-ms time windows with 1-ms steps. Only correct trials were included. For each time window, we performed two partial (Spearman) correlations: 1) between firing rates and choice accuracy while removing the effect of reward expectation, and 2) between firing rates and reward expectation while removing the effect of choice accuracy. Significance was assessed at *p* = 0.05, after multiple comparison corrections using the Benjamini and Hochberg procedure (Benjamini and Hochberg, 1995).

## Acknowledgements

We thank Jean Zweigle for outstanding animal care and training and Michael Suplick for machine shop support (NIH National Eye Institute Core Grant P30 EY001583). We thank other members of Gold and Ding labs for their comments and suggestions. This work was supported by NIH National Eye Institute (R01-EY022411 and R21EY029091; LD and JIG).

## Author contributions

Conceptualization, LD and JIG; methodology, KB and LD; investigation, KB and LD; visualization, LD and JIG; funding acquisition, LD and JIG; project administration, LD; supervision, LD; writing – original draft, LD; writing – review & editing, KB, JIG, and LD.

## Supplemental Figure Legends

**Figure 3-S1:**
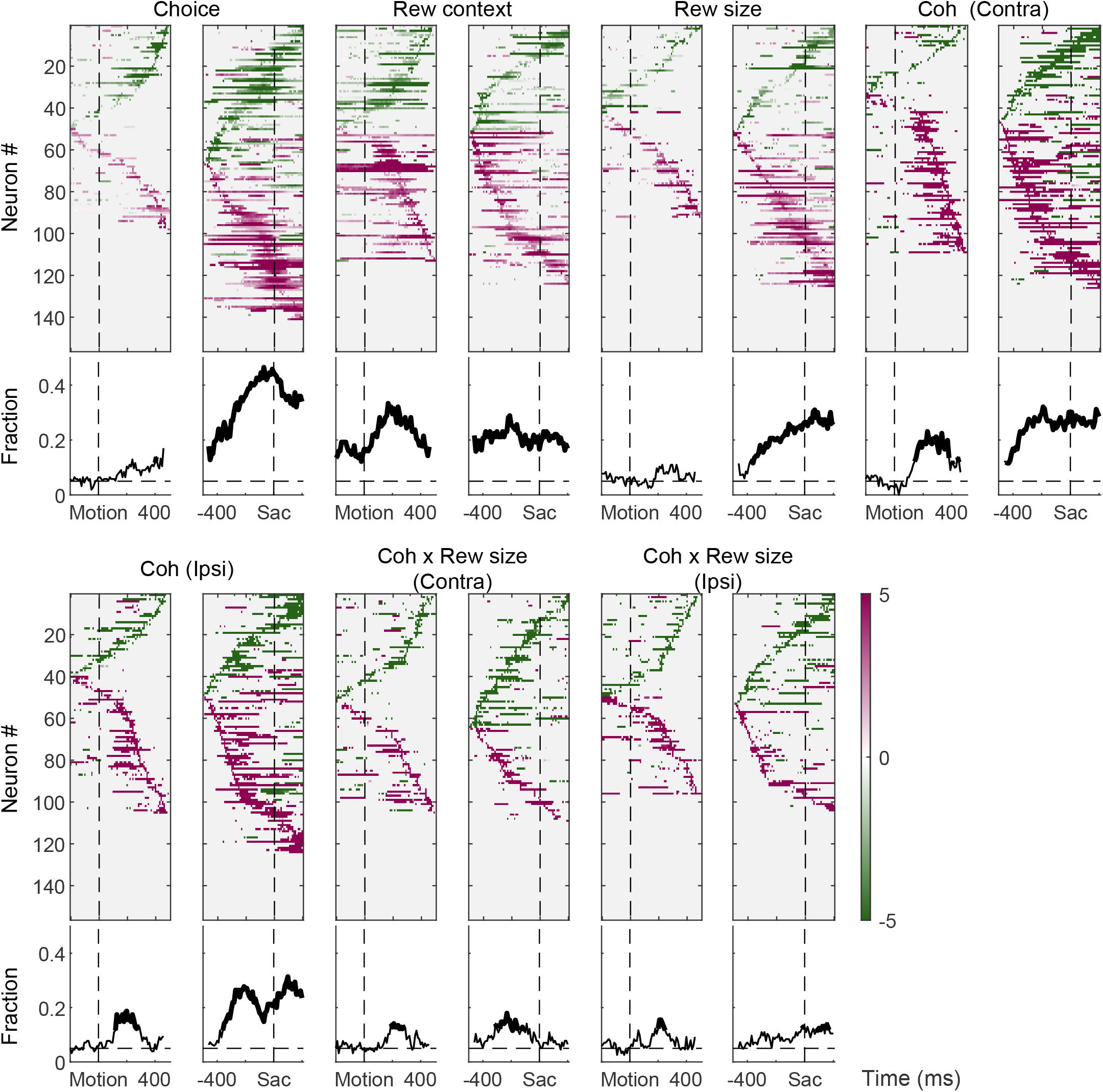
Summary of temporal profiles of modulation of STN activity by decision-related factors. For each regression factor, heatmaps of significant regression coefficients are plotted in the first row for activity aligned to motion (left) and saccade (right) onsets. Neurons are sorted by the timing of peak magnitude of modulation. The fraction of neurons with significant non-zero coefficients are plotted in the second row. Significance for a coefficient was assessed using t-test (p<0.05). For the fraction plots, the dashed horizontal lines represent chance level. Time bins with values that are significantly above chance (chi-square test, p<0.05) are indicated via thicker lines.

**Figure 3-S2:**
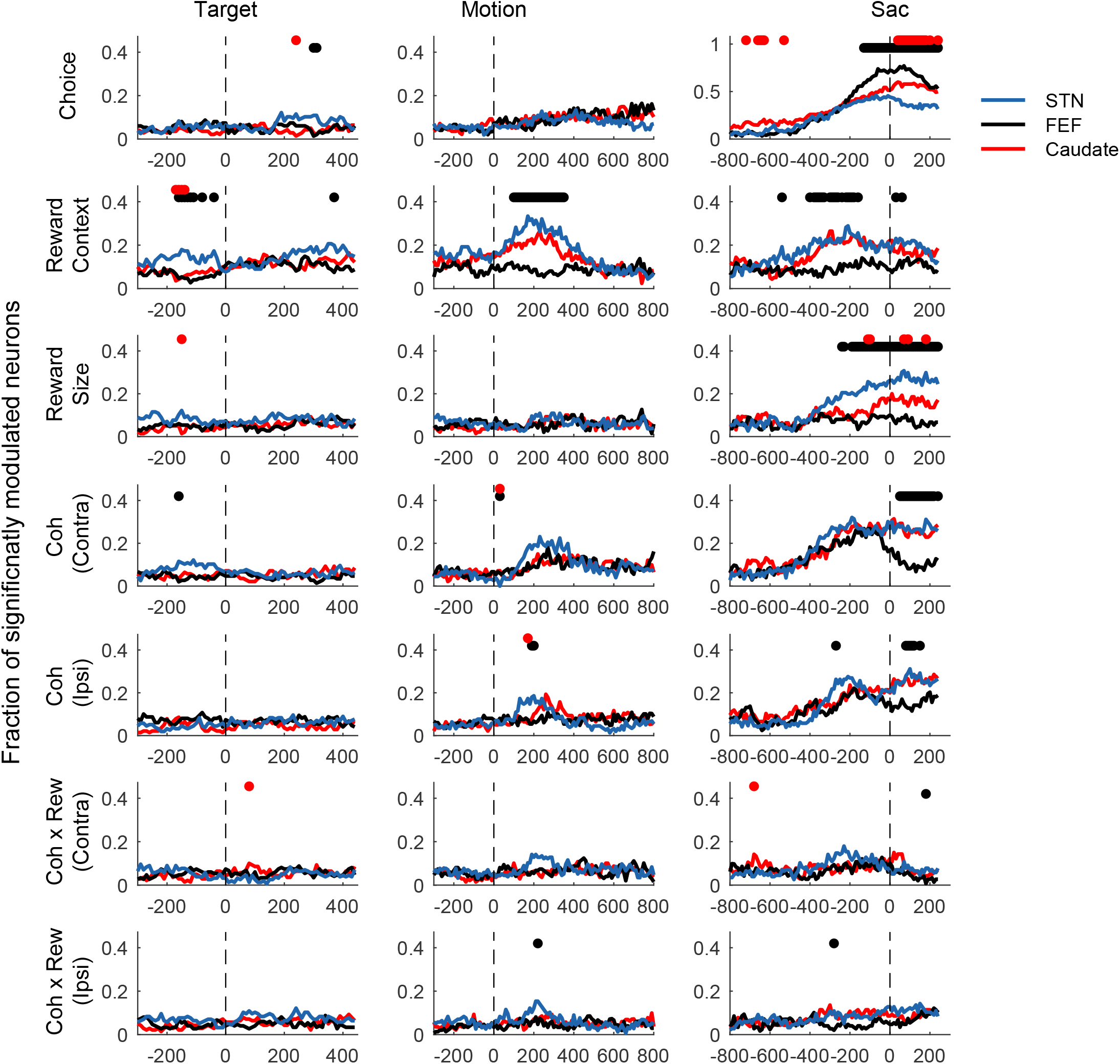
Comparison between STN, FEF, and caudate neurons. Each panel shows the fractions of neurons in the three regions (see legend for colors) with significant coefficients (t-test, p<0.05) for a specific regressor (rows) and activity alignment (columns). Horizontal bars indicate results of chi-square tests using a criterion of p<0.05/3(alignments)/7(regressors). Black horizontal bars indicate significant difference between FEF and STN populations. Red horizontal bars indicate significant difference between caudate and STN populations. FEF and caudate data are from Fan, et al. eLife, 2020.

**Figure 5-S1:**
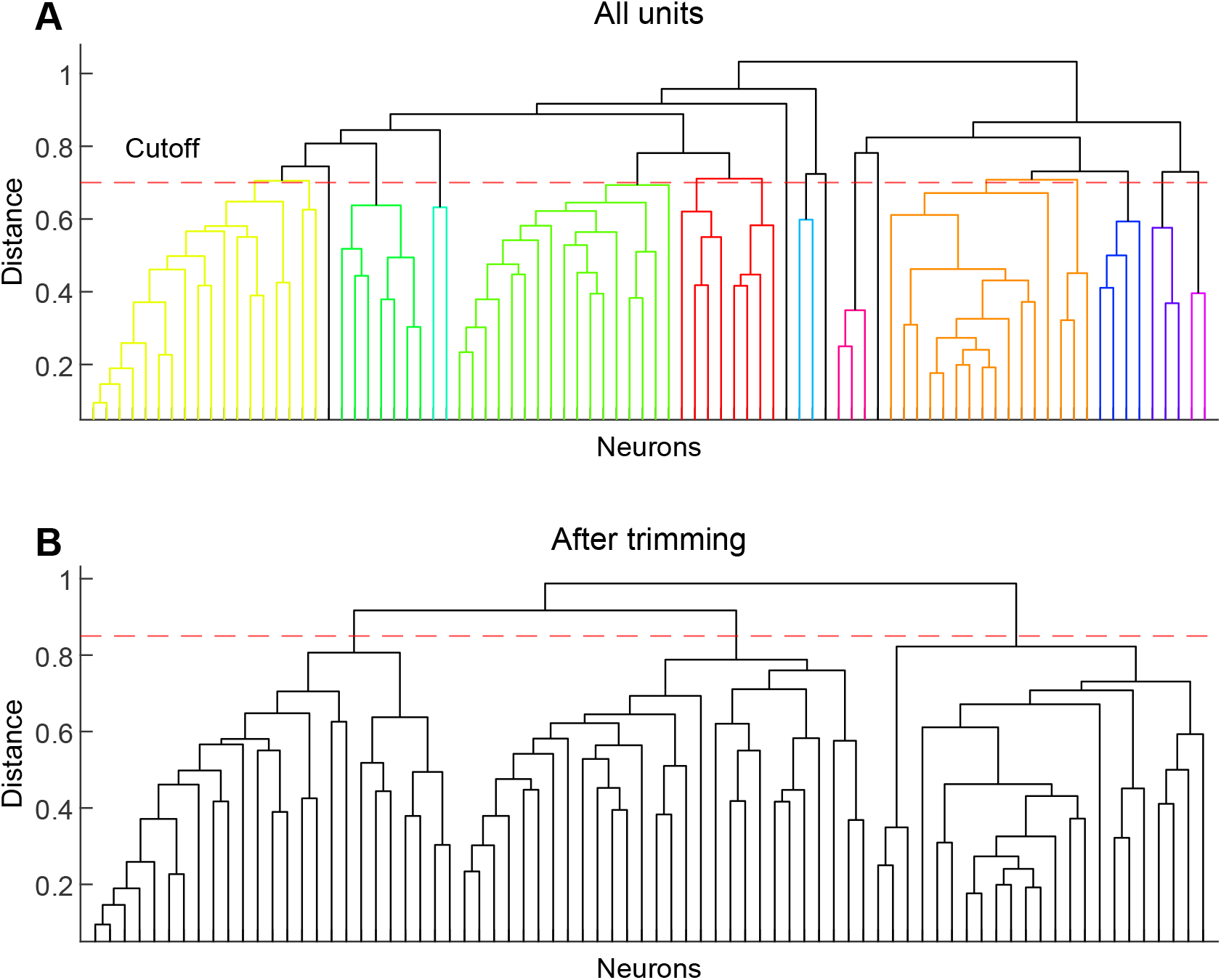
Dendrograms for linkage-based analysis. A, Dendogram computed using all neurons with task-modulated activity (n = 86). A cutoff of 0.7 (red dashed line) was used to identify “outlier” neurons that did not belong to major subpopulations. B, Dendrogram computed using filtered neurons (n = 76). A cutoff of 0.85 was used to cluster the neurons into three groups.

**Figure 5-S2:**
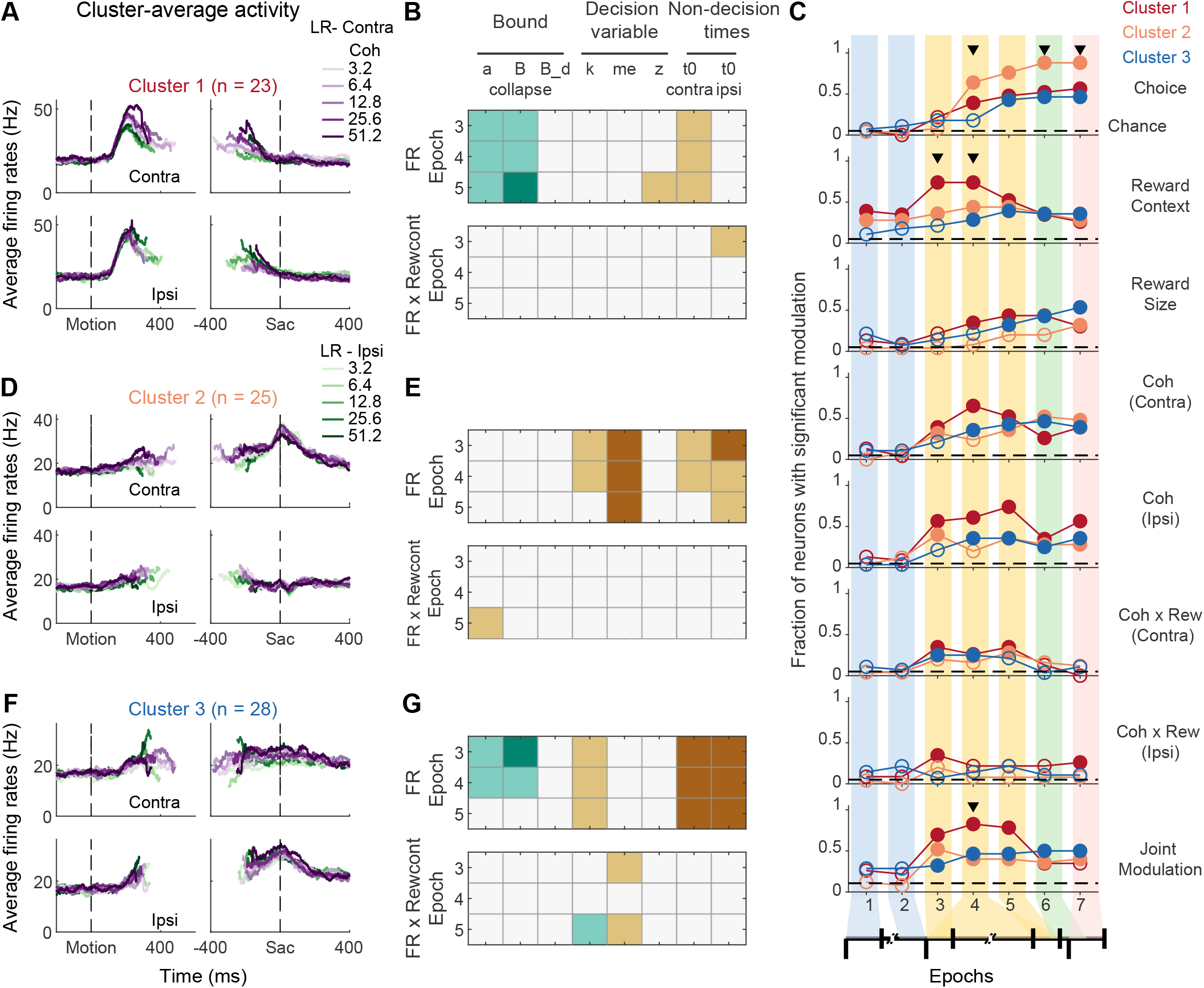
Results from linkage clustering analysis. Same format as Figure 5

**Figure 5-S3:**
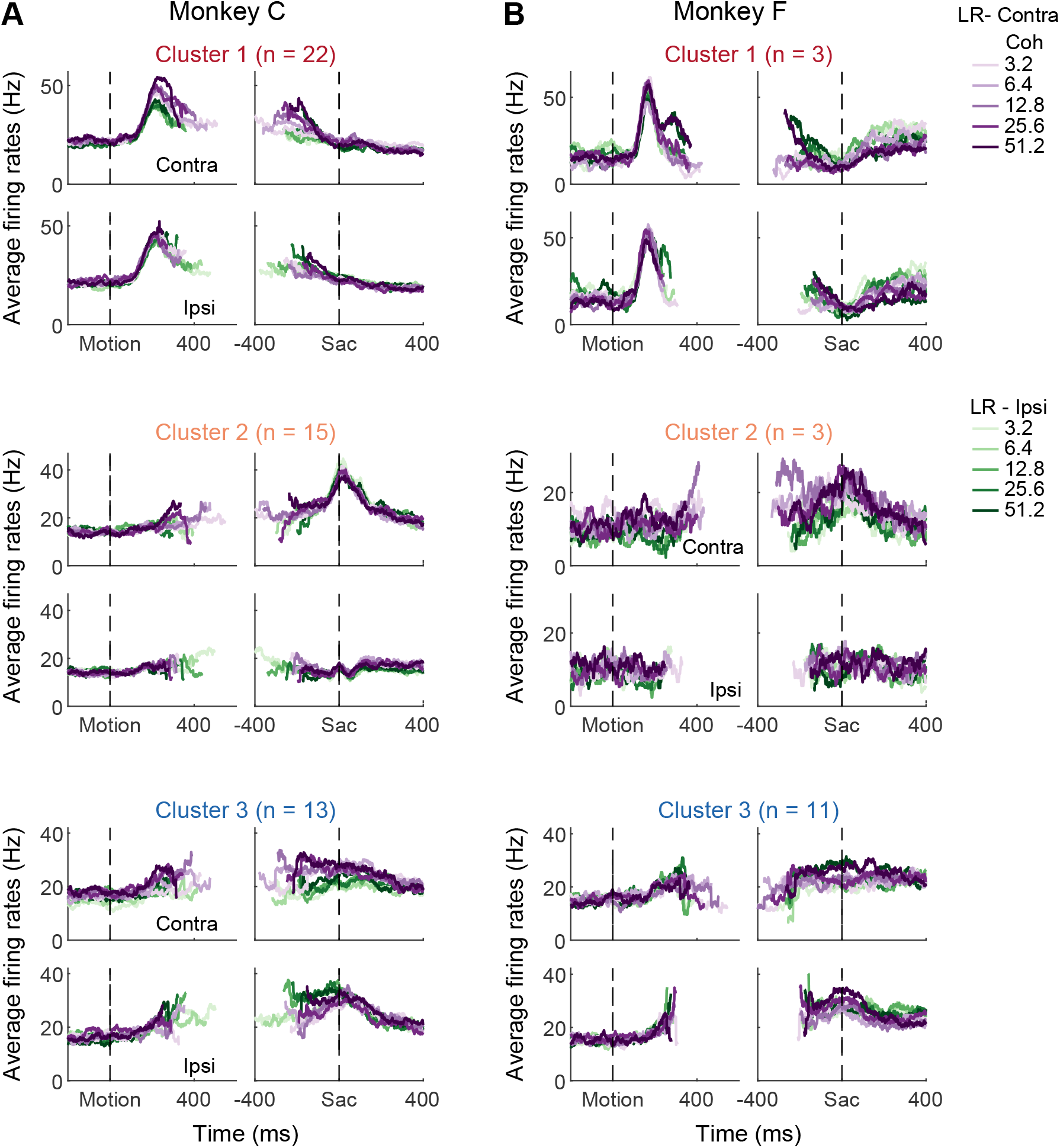
Cluster-average activity for clusters identified using only neurons from monkey C (A) or monkey F (B). Same format as Figure 5A.

**Figure 5-S4:**
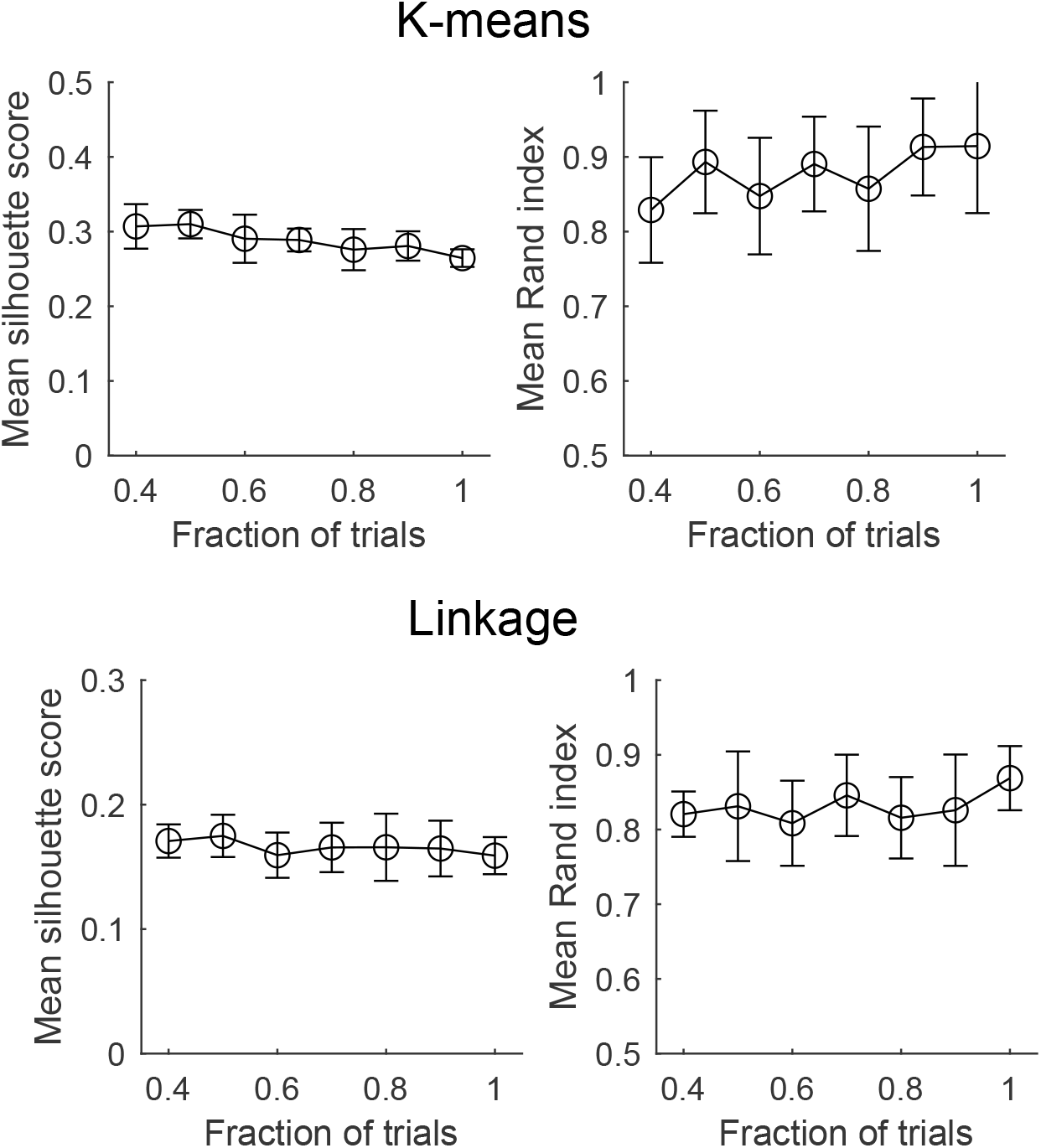
Clustering results are stable with subsets of trials. 10 sets of vectors were generated by sampling with replacement a fraction of trials from each neuron. K-means and linkage clustering were peformed on these resampled vectors. Left column: mean silhouette scores for each fraction value. Right column: mean Rand index between cluster identities of the resampled vectors and the original cluster assignment based on all trials.

**Figure 5-S5:**
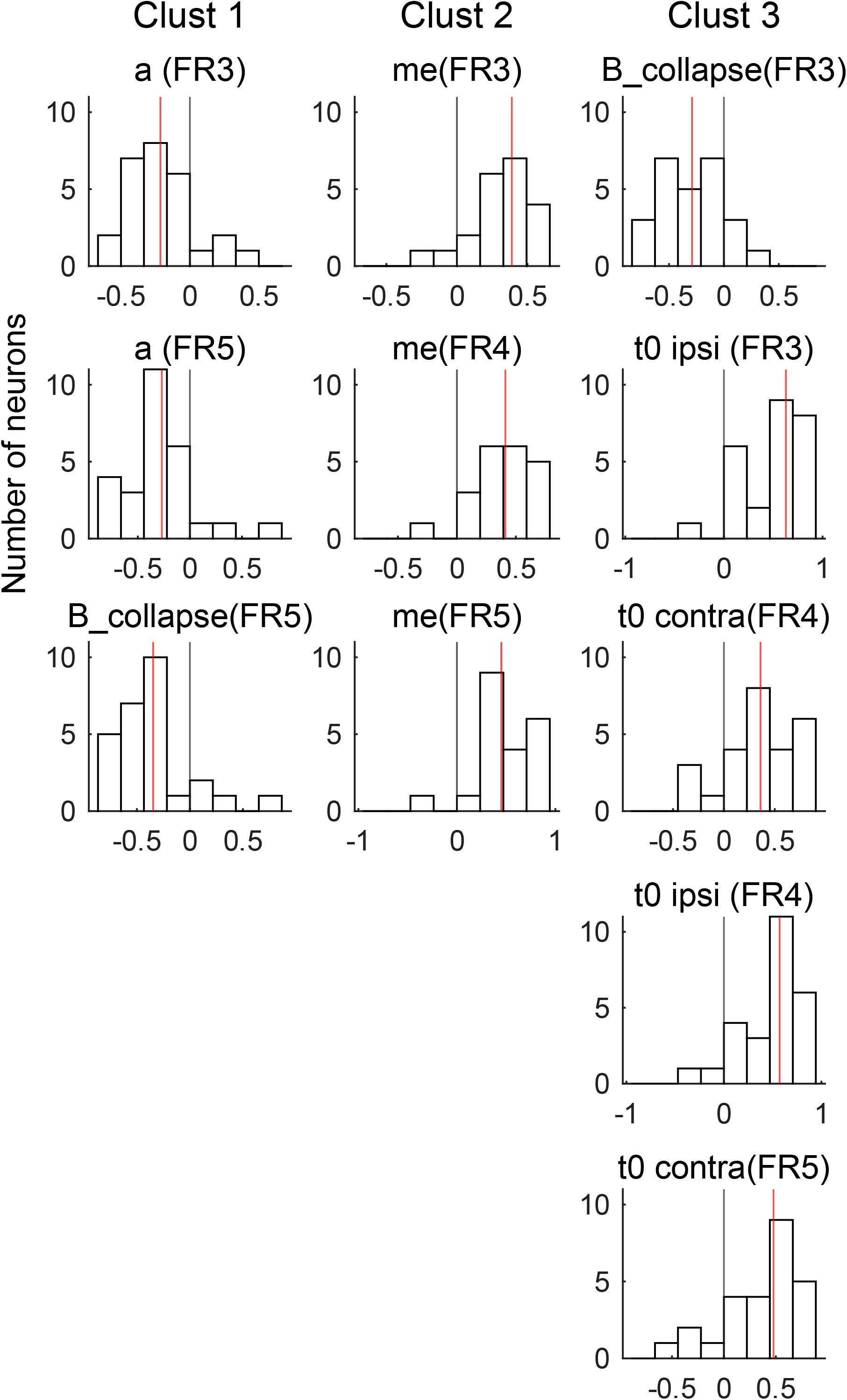
Histograms of significant relationships between neural activity and DDM parameters for the three clusters. Each panel corresponds to a brown/teal box in Figure 5B,E,G. Red lines indicate median values.

**Figure 6S1:**
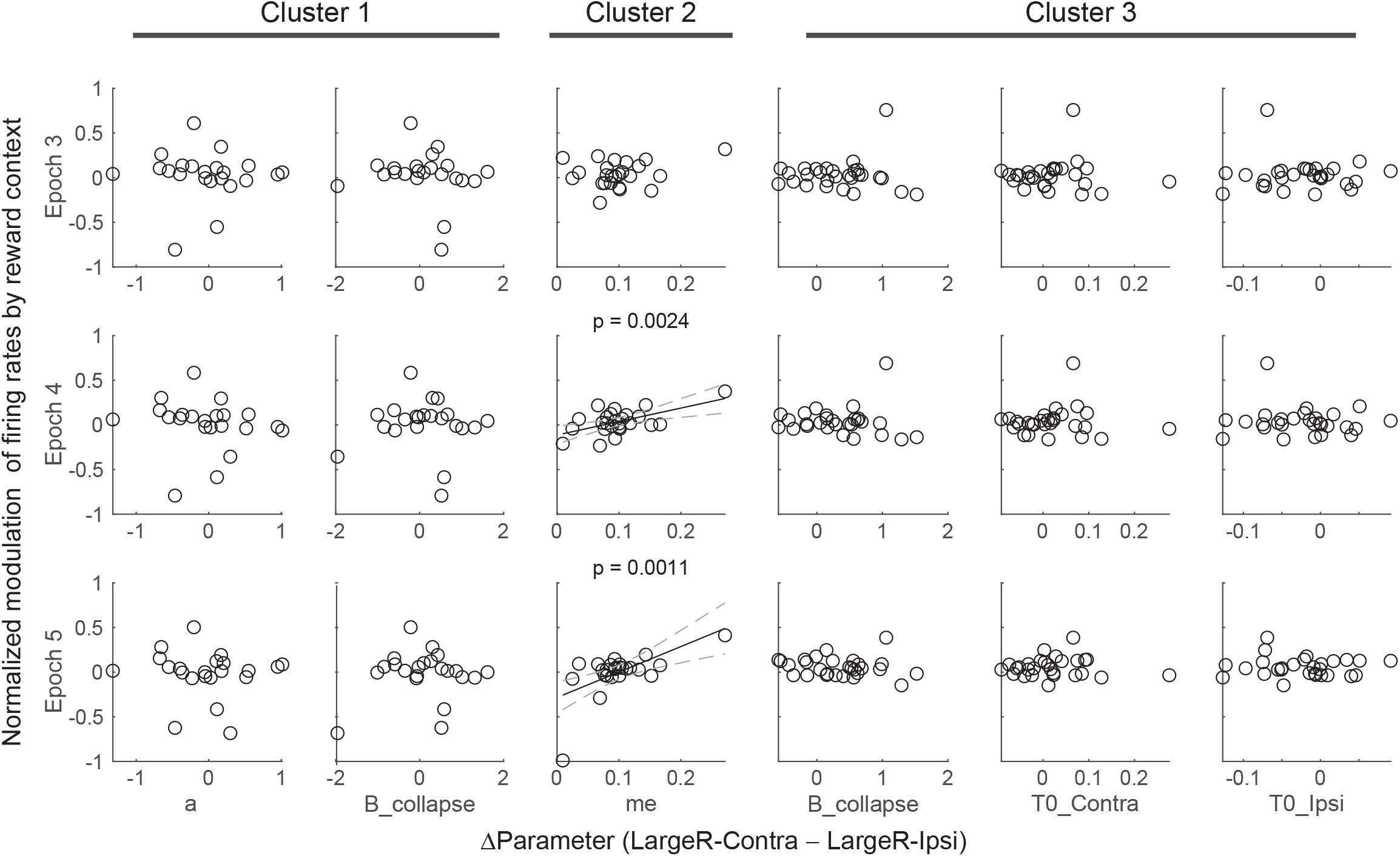
Relationship between reward context modulation of firing rates and reward context modulation of DDM parameters, based on linkage clustering. Same format as Figure 6.

